# Where in the leaf is intercellular CO_2_ (C_i_)? Considerations and recommendations for assessing gaseous diffusion in leaves

**DOI:** 10.1101/2020.05.05.079053

**Authors:** Joseph R. Stinziano, Jun Tominaga, David T. Hanson

**Author notes:** **Corresponding author:** David T. Hanson. Other email addresses.

## Abstract

The assumptions that water vapor exchange occurs exclusively through stomata, that the intercellular airspace is fully saturated with water vapor, and that CO_2_ gradients are negligible between stomata and the intercellular airspace have enabled significant advancements in photosynthetic gas exchange research for nearly 60 years via calculation of intercellular CO_2_ (C_i_). However, available evidence suggests that these assumptions may be overused. Here we review the literature surrounding evidence for and against the assumptions made by Moss & Rawlins (1963). We reinterpret data from the literature by propagating different rates of cuticular water loss, CO_2_ gradients, and unsaturation through the data. We find that in general, when cuticle conductance is less than 1% of stomatal conductance, the assumption that water vapor exchange occurs exclusively through stomata has a marginal effect on gas exchange calculations, but this is not true when cuticle conductance exceeds 5% of stomatal conductance. Our analyses further suggest that CO_2_ and water vapor gradients have stronger impacts at higher stomatal conductance, while cuticle conductance has a greater impact at lower stomatal conductance. Therefore, we recommend directly measuring C_i_ whenever possible, measuring apoplastic water potentials to estimate humidity inside the leaf, and exercising caution when interpreting data under conditions of high temperature and/or low stomatal conductance, and when a species is known to have high cuticular conductance.

**Highlight:** Leaf water vapor and CO_2_ exchange have been successfully used to model photosynthetic biochemistry. We review critical assumptions in these models and make recommendations about which need to be re-assessed.

## Introduction

Nearly 60 years ago, Moss and Rawlins (1963) introduced a calculation to estimate intercellular CO_2_ concentrations (C_i_) under the assumption that all water flux out of the leaf occurs through the stomata. The importance of their findings was such that it quickly became dogma and is severely under-cited (<100 citations according to Google Scholar) despite underlying nearly every measurement of stomatal conductance (g_s_) and C_i_. C_i_ now plays a central role in plant gas exchange research: it is used to derive parameters for a biochemical model of leaf gas exchange through measurements of net CO_2_ assimilation (A_net_) responses to CO_2_ (Farquhar *et al*., 1980), which in turn can be used to drive photosynthesis in coupled vegetation-climate models (e.g. Oleson *et al*., 2013), and C_i_ is a necessary starting point for estimating CO_2_ fluxes within leaves all the way to the site of carboxylation (Evans *et al*., 1986). However, this approach uses myriad assumptions that are generally not realistic (and almost physically impossible in other cases), including saturated vapor pressure in the leaf (Hygen, 1951, 1953; Slavik, 1958; Jarvis & Slatyer, 1970; Ward & Bunce, 1986; Egorov & Karpushkin, 1988; Karpushkin, 1994; Campbell & Norman, 1998; Canny & Huang, 2006; Cernusak et al., 2018), negligible cuticular conductance (Boyer et al., 1997; Meyer & Genty, 1998; Šantrůček et al., 2004; Boyer 2015a; Tominaga & Kawamitsu, 2015a; Tominaga et al., 2018), homogenous stomatal conductance (Downton et al., 1988; Terashima et al., 1988; Buckley et al., 1997; Meyer & Genty, 1998), no CO_2_ gradients within leaves (i.e. infinite CO_2_ conductance; Parkhurst, 1984; Long et al., 1989), no air pressure gradients within the leaf (Leuning, 1983; Dacey, 1987), strict Fickian diffusion of CO_2_ into leaves (Leuning, 1983; Dacey, 1987), and resistances to gas diffusion are additive (i.e. follow the Ohm’s law analogy, Parkhurst, 1984). For the purposes of this review, we define infinite conductance as a conductance high enough that a negligible concentration gradient forms between the two compartments in question. Here we review the concept of C_i_, g_s_, and assumptions inherent in their calculations, and provide recommendations on moving beyond the Moss and Rawlins (1963) paradigm of assumptions.

### The Moss and Rawlins (1963) paradigm

To calculate stomatal resistance, Gaastra (1959), with Brown and Escombe’s (1900) Ohm’s law analogy of resistances to gas diffusion (although this may not necessarily hold true: see Parkhurst, 1984), introduced a series of Equations: 

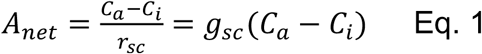

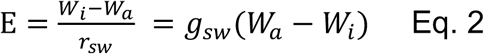

where *A*_*net*_ is net CO_2_ assimilation, *C*_*a*_ is the CO_2_ concentration external to the leaf, *r*_*sc*_ and *g*_*sc*_ is the stomatal resistance and conductance (reciprocal of resistance) to CO_2_, *E* is transpiration through the stomata, *W*_*a*_ and *W*_*i*_ are the water vapor concentration external to the leaf and in the intercellular airspace, *r*_*sc*_ and *g*_*sc*_ is the stomatal resistance and conductance to water vapor. Considering the boundary resistance was negligibly small in the measurement system (which we also assume through this review), Moss and Rawlins (1963) expressed the diffusion properties of CO_2_ and water vapor in leaves as: 

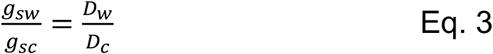

where *D*_*w*_ and *D*_*c*_ are the diffusion coefficients for CO_2_ and water vapor in air. *D*_*w*_/*D*_*c*_ is usually assumed to be equal to ∼1.6 (Li-Cor, 2019; although see Holmgren et al 1965 Physiologia Plant. 18:557 where they use 1.7). Note that 1) Massman (1998) reported a mean ratio of 1.577 with uncertainties as high as 7% for the diffusivity of water vapour, 2) the number is valid for Fickian diffusion, and 3) this may vary if the stomatal pore size is small enough that Knudsen diffusion occurs instead of Fickian diffusion (e.g. Parkhurst, 1994). Solving the system of Eqs.1-3 they obtained: 

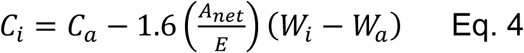

where *W*_*i*_ is assumed to be saturated for the leaf temperature (*T*_*leaf*_). Using conductance term, Eq. 4 can be rewritten as: 

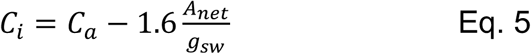

Given that *C*_*a*_ and *A*_*net*_ are directly measurable, *g*_*sw*_, calculated from *E* and water vapor gradients (*W*_*a*_-*W*_*i*_) (Eq. 2), would be the critical parameter that determines validity of *C*_*i*_ (though measurement precision of *C*_*a*_ and *A*_*net*_ affects the calculation).

Based on the above, Moss and Rawlins (1963) introduced assumptions (both explicitly and implicitly) into the calculation of *C*_*i*_:

1. All transpired water flows through stomata
2. No CO_2_ or H_2_O gradients within leaves (internal conductance, g_ias_ = ∞)
3. *W*_*i*_ is at saturation (W_i_ = saturation vapor pressure, e_s_; mesophyll apoplast water potential, Ψ_m,apo_ = 0)
4. Uniform stomatal apertures over leaf surfaces
5. No air pressure gradients across leaf surfaces
6. 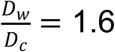 (based on Fickian diffusion in air)
7. One-dimensional models approximate the three-dimensional leaf

For the purposes of this review, we are going to address the data surrounding the implications and failures of assumptions 1 to 3 in relation to *C*_*i*_ calculations and data derived from *C*_*i*_ and give a cursory overview on the remaining assumptions. Note that these assumptions equally apply to the von Caemmerer and Farquhar (1981) modification to Moss and Rawlins (1963).

#### What do tests of Moss and Rawlins (1963) say?

The first test of the calculation was done in 1982 (Sharkey *et al*., 1982) whereby *C*_*i*_ was directly measured and compared to the calculated value based on the Moss and Rawlins assumption. Sharkey *et al*. (1982) used a dual open-flow/closed-flow system whereby one side of the leaf was equilibrated in a closed system to measure *C*_*i*_, while the other side was measured with the open-flow system used to calculate *C*_*i*_. Note that this method requires a steady-state setup and assumes that the CO_2_ in the closed-flow system is in equilibrium with the internal airspace of the leaf. Sharkey *et al*. (1982) concluded that the calculated values were in “good agreement” with the measured values, although there were some deviations less than ±20 μmol mol^-1^ (Table 1). They also reported that at very high vapor pressure difference (VPD) and conductance less than 60 mmol m^-2^ s^-1^, the calculated *C*_*i*_ increased while the measured *C*_*i*_ decreased. Since then, fewer than 10 studies have assessed the Moss and Rawlins (1963) assumptions (Table 1) despite the fact that it is the foundation for a broad range of research activities. These studies employed direct *C*_*i*_ measurements similar to Sharkey et al. (1982), except for Boyer et al. (1997). Intriguingly, most evidence suggests that there are issues in the assumptions that lead to discrepancies between measured and calculated *C*_*i*_ (Table 1), with explanations ranging from cuticular water loss (Boyer *et al*., 1997), to patchy stomatal apertures (Downton et al., 1988; Terashima et al., 1988; Meyer & Genty, 1998) and intra-leaf CO_2_ gradients (Sharkey et al., 1982; Mott & O’Leary, 1984; Parkhurst *et al*., 1988; Parkhurst & Mott, 1990; Parkhurst, 1994).

**Table 1.**
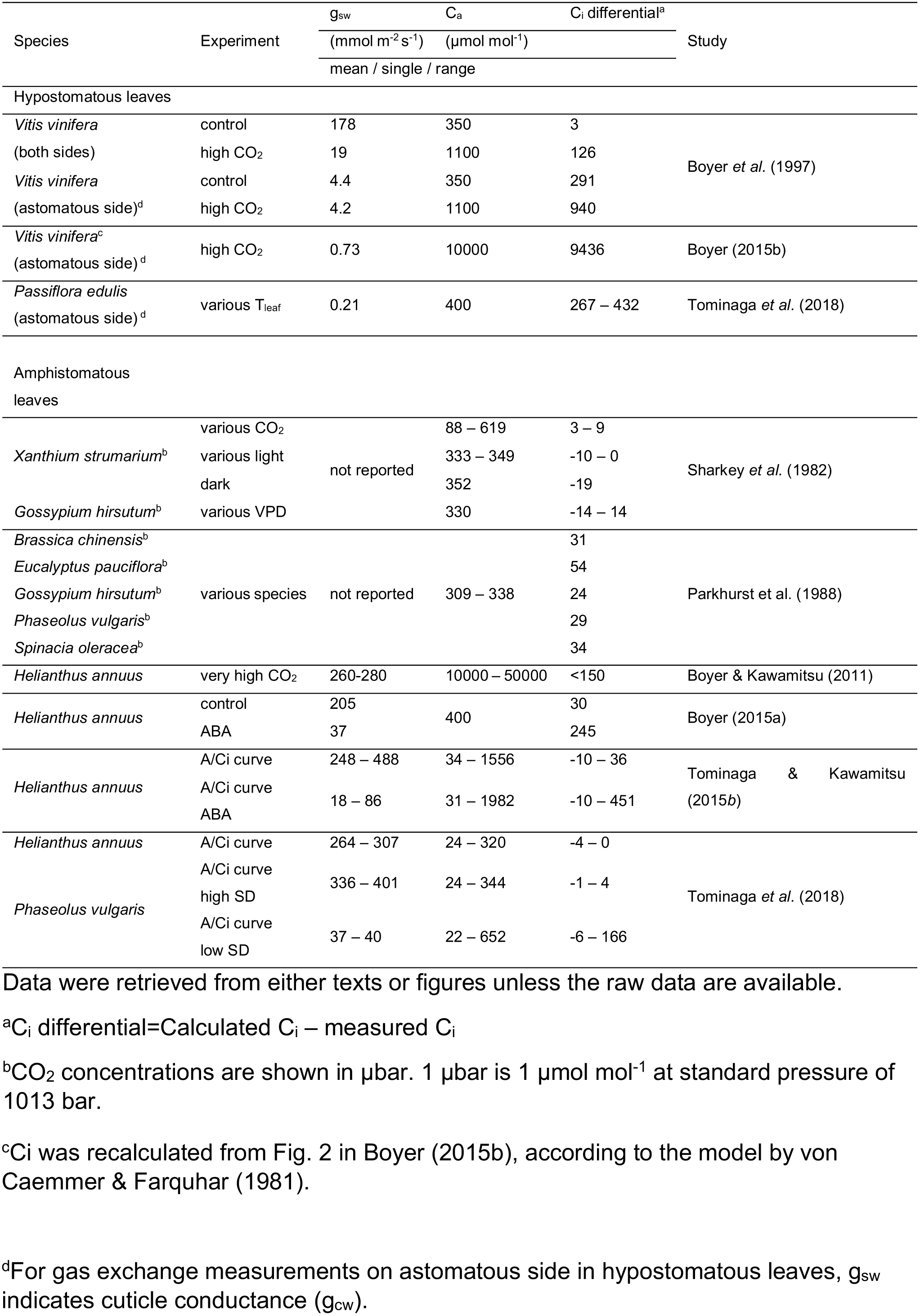
Reported *C*_*i*_ for direct tests of the Moss and Rawlins (1963) assumptions.

#### Assumption 1 – all water flows through stomata

Cuticular water loss is a proposed explanation for the reported discrepancies (Kirschbaum and Pearcy, 1988; Boyer *et al*., 1997; Meyer & Genty, 1998; Boyer, 2015a; Tominaga & Kawamitsu, 2015a, Tominaga et al., 2018). Using hypostomatous leaves of grape (*Vitis vinifera* L.), Boyer et al. (1997) measured gas exchange through the adaxial stomata-free side while the abaxial stomatous side was sealed. In this circumstance, *C*_*i*_ was estimated to be near the compensation point (50 μmol mol^-1^) with a little CO_2_ flux on the cuticular side while calculations showed *C*_*i*_ close to *C*_*a*_. As a result, the calculated *C*_*i*_ was larger than the actual value by over a hundred μmol mol^-1^ (astomatous side of *Vitis vinifera* in Table 1). Later, this result was reproduced in the same species (Boyer 2015b) with a direct system that reached *C*_*a*_ as high as several % to detect very small CO_2_ fluxes through the cuticle (Boyer & Kawamitsu, 2011), as well as in passion fruits (*Passiflora edulis* Sims) (Tominaga et al., 2018). Boyer (2015b) found that the measured *C*_*i*_ increased by only 2 μmol mol^-1^ above the CO_2_ compensation point of 44 μmol mol^-1^ despite the large CO_2_ gradients (10000 μmol mol^-1^) across the cuticular surface, suggesting that the cuticle is an effective barrier against CO_2_ diffusion. Transpiration was always greater than assimilation through the cuticle, causing cuticle conductance for water vapor (*g*_*cw*_) to be 20-40× larger than cuticle conductance for CO_2_ (*g*_*cc*_), which is a much higher ratio than the 1.6 assumed for stomata (Eq. 3). This is likely because the pathway for CO_2_ in these experiments was from the leaf airspaces through the epidermal cells and cuticle, whereas the pathway for water diffusion was from the epidermal surface through the cuticle.

Calculations (Eq. 4) consistently overestimate the *C*_*i*_ for the astomatous side of hypostomatous leaves (Table 1). In contrast, *C*_*i*_ values were in closer agreement between direct measurements and calculations in the high CO_2_ tests for stomatous leaf surfaces in sunflower (*Helianthus annuus* L.) (Boyer & Kawamitsu (2011) in Table 1). It should be noted that von Caemmerer and Farquhar (1981) slightly modified Eq. 4 by including a ternary effect term that describes the hinderance of CO_2_ diffusion into the leaf due to the much larger flux of H_2_O out of the leaf (Eq. B18 in von Caemmerer & Farquhar, 1981), and this version is used more generally and also in Table 1. Boyer & Kawamitsu (2011) experimentally validated this modification under high *C*_*a*_ that enhanced the ternary effect, and thus direct measurements include this effect.

To see the cuticle effect on both leaf sides (cuticle plus stomata), Boyer (1997) recalculated *C*_*i*_ in the standard measurements for both sides using the cuticle conductance determined on the same leaf (*Vitis vinifera* (both sides) in Table 1). The results suggest that the cuticle effect can be substantial—*C*_*i*_ differential is 126 μmol mol^-^ 1 at 1100 μmol mol^-1^ *C*_*a*_— when *g*_*sw*_ is relatively small, but only marginal—*C*_*i*_ differential of 3 μmol mol^-1^ at 350 μmol mol^-1^ *C*_*a*_— when *g*_*sw*_ is relatively large. This conclusion was recently confirmed with direct measurements in amphistomatous sunflower leaves with stomata closed by feeding ABA (Boyer, 2015a; Tominaga & Kawamitsu, 2015a), and amphistomatous bean (*Phaseolus vulgaris* L.) leaves with low stomatal density (SD) (Tominaga et al., 2018), as summarized in Table 1. In these studies (Tominaga & Kawamitsu, 2015a; Tominaga et al., 2018), calculation and direct measurements draw essentially the same *A*/*C*_*i*_ response curves when *g*_*sw*_ was large (*g*_*sw*_>250 mmol m^-2^ s^-1^) with open stomata and/or high SD. In contrast, when *g*_*sw*_ was small (*g*_*sw*_<50 mmol m^-2^ s^-^1) with closed stomata and/or low SD, *A*/*C*_*i*_ curves were depressed due to over-estimation of the calculated *C*_*i*_. The similar depression was also confirmed with the standard open-flow measurements for both sides (Tominaga & Kawamitsu, 2015a), as was observed previously in similar ABA treatments (Downton et al., 1988; Terashima et al., 1988). Clearly, this should create a problem for interpreting gas exchange measurements. Cuticular water loss also causes calculated *C*_*i*_ to be lower than the actual value when CO_2_ is diffusing out from the leaf as it overestimates the CO_2_ transfer through stomata regardless of diffusional direction. In accordance, negative *C*_*i*_ differentials were found with negative *A*_*net*_ in dark, and low *C*_*a*_ in light (Table 1).

There are debates as to whether and when cuticular water loss would be a significant portion of water loss across the leaf (Ledford, 2017). Generally, we would expect cuticle conductance to be more significant at low values of g_sw_ as noted above (Meyer and Genty, 1998; Flexas *et al*., 2002; Lawlor, 2002) and under heat stress where the cuticle could undergo a state change to become very permeable to water (although note that cuticular melting may not occur until temperatures >60 °C, Bargel et al., 2006). But how large is leaf cuticle conductance and water loss? Unfortunately, biologists studying cuticle properties often focus on cuticle permeance (units: m s^-1^), which can hinder comparisons with gas exchange where conductance and flux are normally measured (units mol m^-2^ s^-1^). A series of equations permits the calculation of conductance and flux from permeance. For conductance (Hall, 1982; Nobel, 1991): 

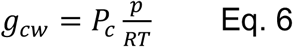

Where g_cw_ is cuticular conductance for water vapor (mol m^-2^ s^-1^), P_c_ is cuticular permeance (m s^-1^), p is atmospheric pressure (Pa), R is the universal gas constant (8.314 m^3^ Pa K^-1^ mol^-1^), and T is temperature (K). And for the flow rate of water across the cuticle (flux, transpiration), assuming steady-state conditions (Riederer and Schreiber, 2001): 

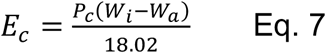

Where E_c_ is cuticular transpiration (water flux) across the cuticle (mol m^-2^ s^-1^), P_c_ is cuticular permeance (m s^-1^), W_i_ is the water vapor concentration adjacent to the outer epidermal wall (g m^-3^), W_a_ is the water vapor concentration at the leaf surface (g m^-3^), and 18.02 is the molar mass of water (g mol^-1^).

Cuticle permeances (m s^-1^) available in Riederer & Schreiber (2001) were converted into g_cw_ and E_c_ values wherever sufficient data were available in the original papers to perform the calculations (Table 2). Mean g_cw_ calculated from Riederer and Schreiber (2001) was 0.511 ± 0.101 mmol m^-2^ s^-1^ (range: 0.015 to 5.862 mmol m^-2^ s^-1^), while mean E_c_ was 15.18 ± 2.66 μmol m^-2^ s^-1^ (range: 0.46 to 134.36 μmol m^-2^ s^-1^) (Table 2; Fig 1a). How do these g_cw_ values compare to stomatal conductance?

**Table 2.**
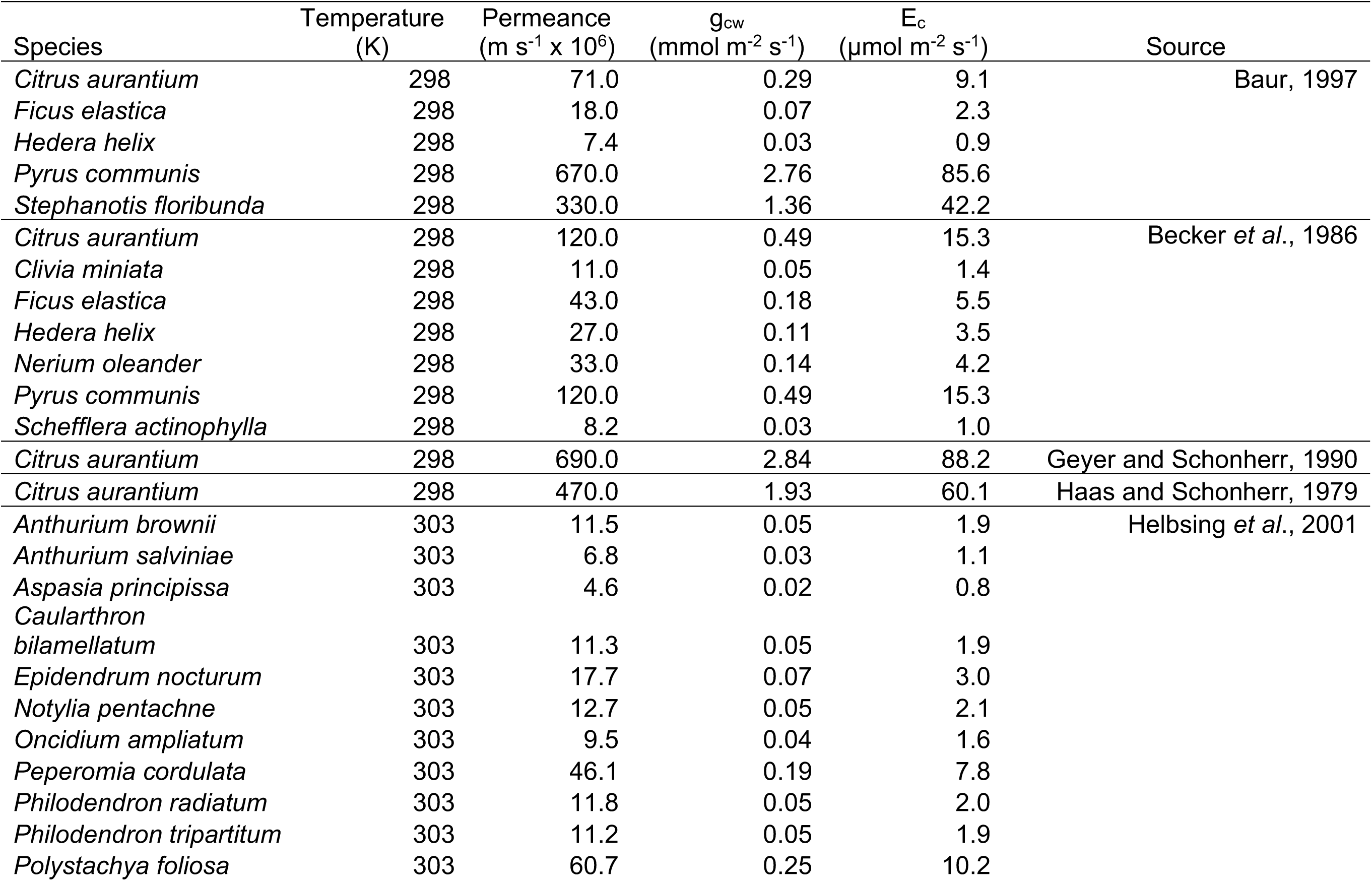
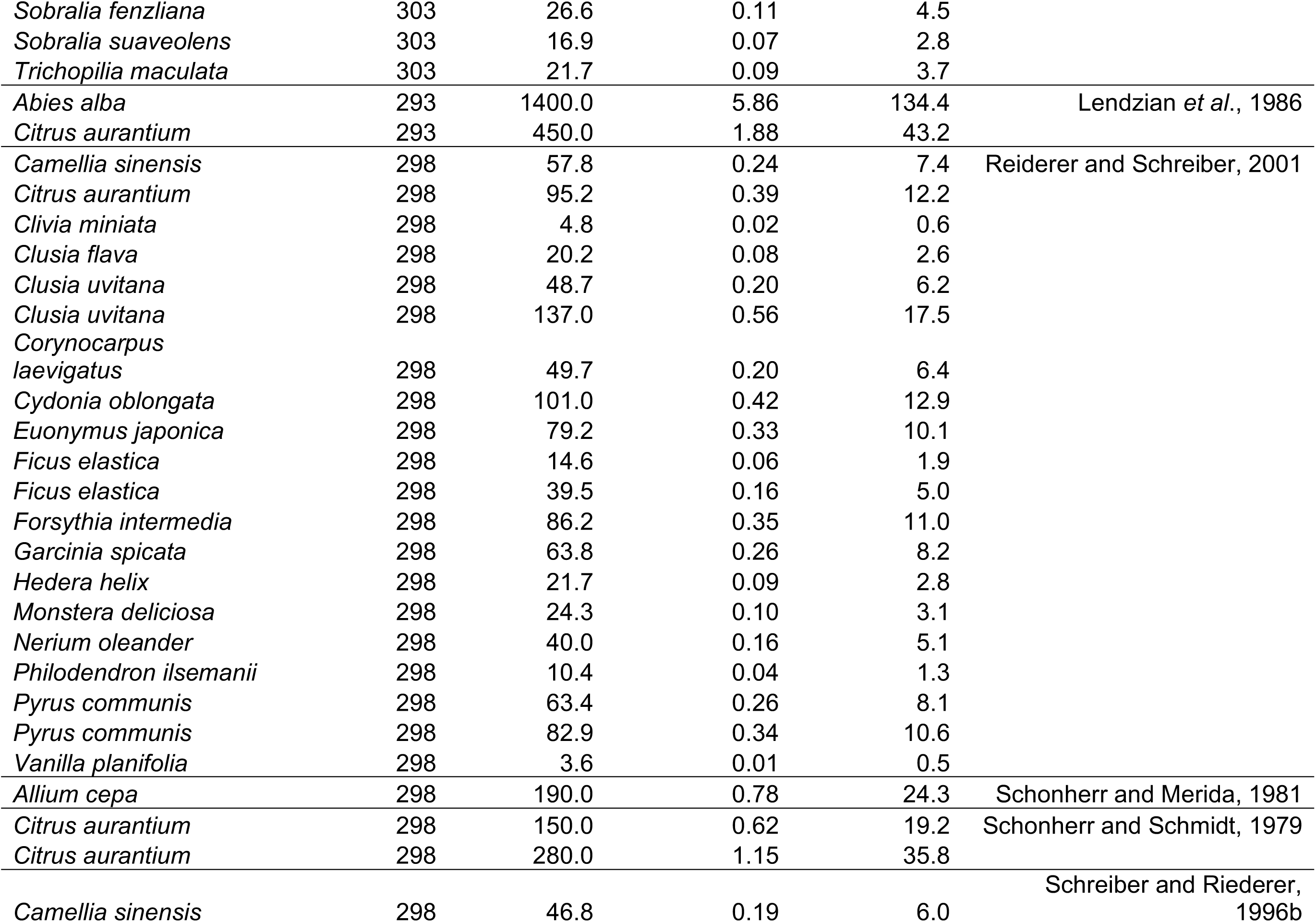
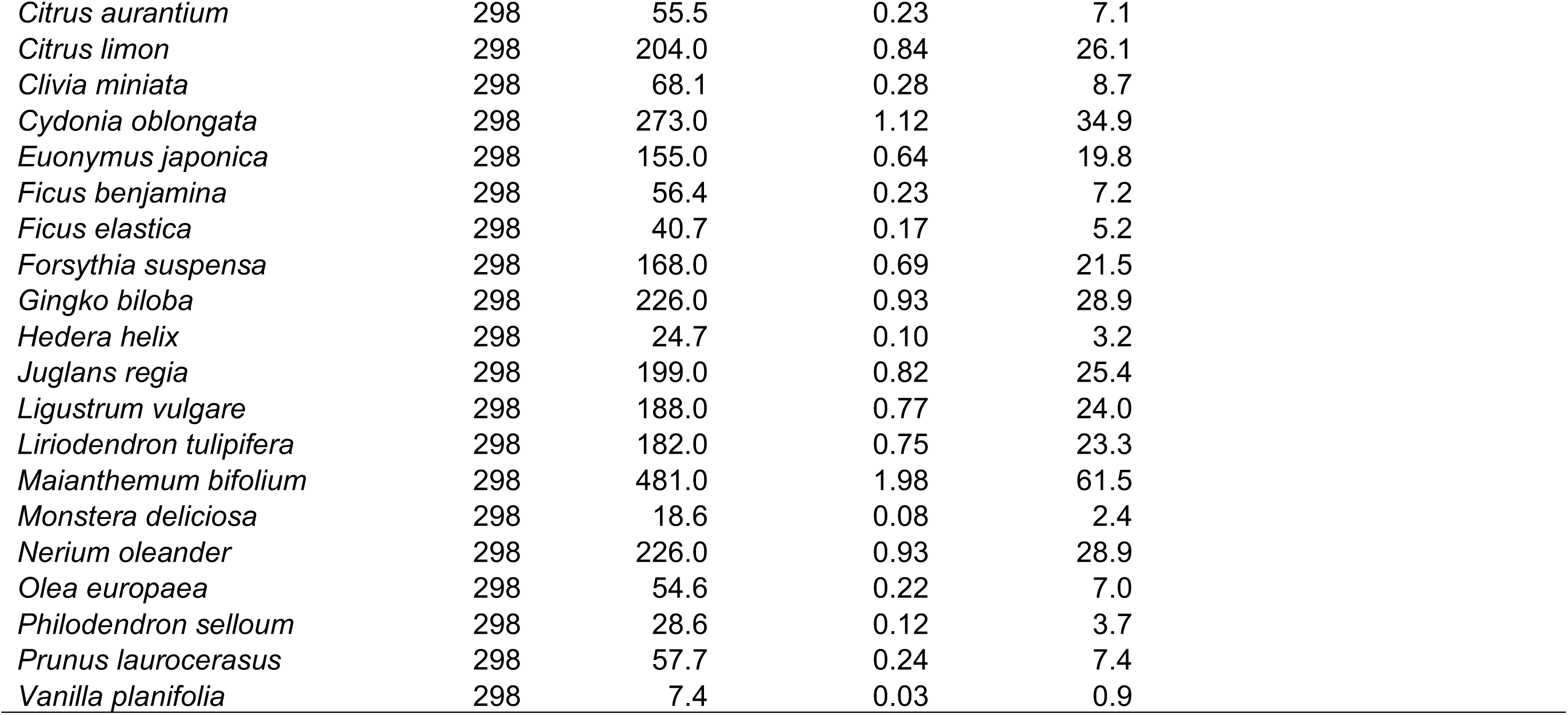
Cuticular permeances from Riederer and Schreiber (2001) re-interpreted as cuticular conductance (g_cw_) and transpiration (E_c_).

**Figure 1.**
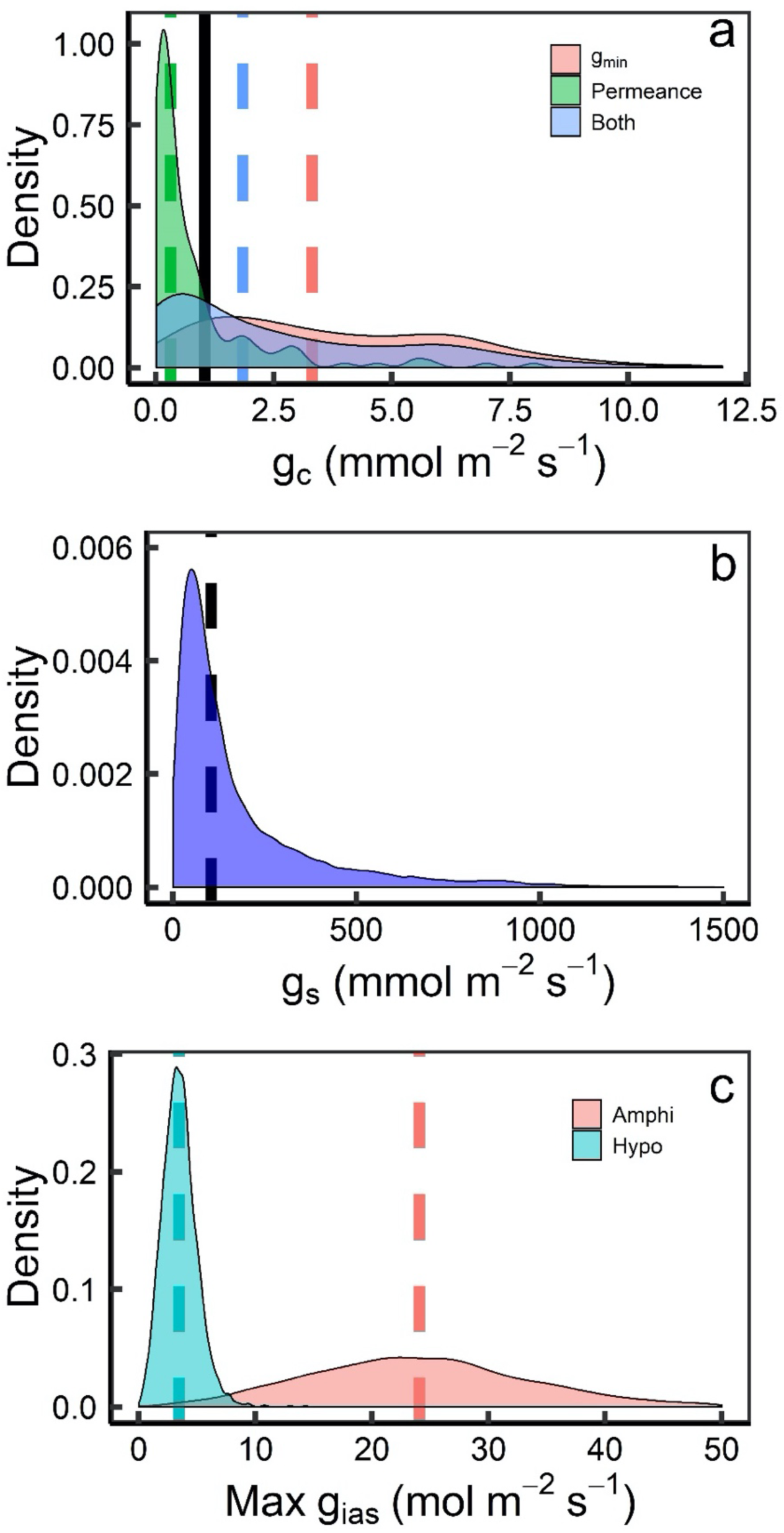
Density plot of measured (a) cuticle (g_c_), (b) stomatal (g_s_) values, and (c) maximum theoretical intercellular airspace conductance (g_ias_) values assuming diffusion through 12.5% of leaf thickness for amphistomatous leaves (Amphi) and 87.5% of leaf thickness for hypostomatous leaves (Hypo). (a) Data was compiled from Riederer and Schreiber (2001) and the supplementary information from Schuster et al. (2017). Black line indicates 1% of the median value from (b), while dashed lines indicate method-specific medians. (b) Data was compiled from Lin et al. (2015) and Smith and Dukes (2017). (c) Leaf thickness data used to calculate maximum g_ias_ from Onoda et al. (2011).

We compared the range in g_cw_ values above to the stomatal conductances (g_sw_) reported in Douthe *et al*. (2011), Vrábl *et al*. (2009), and Scafaro *et al*. (2011) (which we also use to test the implications of g_cw_ on mesophyll conductance, g_m_, below; see Pons et al., 2009 for a discussion of this). Given that measured g_sw_ from the studies used for the g_m_ analysis varied from 43.6 to 1,253 mmol m^-2^ s^-1^ (mean: 474 ± 44 mmol m^-2^ s^-1^), a quick estimate suggests that g_cw_ could range between 0.001 and 13% of g_sw_ measurements (calculation based on means: 0.1% of g_sw_), which may have significant implications for gas exchange measurements in certain species under some conditions.

Analyzing Eq. 7, there are two unknowns (P_c_ and W_i_, though P_c_ is measurable in astomatous cuticles), necessitating an assumption about the value of either P_c_ or W_i_. Typically, W_i_ is assumed to be equal to the water vapor concentration of the saturation vapor pressure at T_leaf_. However, it is difficult to assess whether this assumption holds, as the site of evaporation for cuticle conductance is within the cell wall of the epidermis and reflects a different site of evaporation than for airspace W_i_. This assumption may be broken during the dry-down experiments for gravimetrically-determined cuticular water loss. During these measurements, water loss rates decline over time until a breakpoint and a constant rate of water loss are achieved. Beyond the breakpoint, stomata are assumed to be closed. Furthermore, despite the leaf having lost a substantial amount of water, the W_i_ assumption is used to calculated P_c_, with W_i_ assumed to be the same for the cell wall of the epidermis and the intercellular airspace. Therefore, at a given value of E_c_, when the W_i_ assumption is violated, then P_c_ will change. In this way, it is possible that much of the literature on cuticular water loss is mis-estimating cuticular permeances, and therefore cuticular conductance. This could explain why gas exchange estimates of cuticular conductance often far exceed the conductance measured using the gravimetric method or isolated cuticles, although leaky stomata in the gas exchange methods could contribute to these differences as well.

#### Assumption 2 – no CO_2_ or H_2_O gradients within leaves

Due to finite intercellular CO_2_ conductance in leaves (*g*_*ias*_), adaxial-abaxial gradients of *C*_*i*_ must exist along the mesophyll cells (CO_2_ sink). In amphistomatous leaves, CO_2_ diffuses through stomata on both sides, and the diffusion path meets somewhere in the middle of the leaf where the gradient ends. Because CO_2_ diffuses slowly through cuticle, larger *C*_*i*_ gradients would develop in hypostomatous leaves than in amphistomatous leaves as the path-length could be longer in the airspace (Parkhurst and Mott, 1990; Evans and Loreto, 2000). Direct measurement technically alters amphi-to hypostomatous leaves by closing one surface, thereby doubling the diffusion path (e.g., Fig. 9 in Boyer & Kawamitsu, 2011). While CO_2_ is entering through one side, direct measurements measure the CO_2_ equilibrated at the opposite side—end of the diffusion path—and thus measures the lowest *C*_*i*_ for the gradient. Therefore, positive *C*_*i*_ differentials observed in amphistomatous leaves may be associated with the gradient. Parkhurst et al. (1988) explored this effect by observing 20-60 μmol mol^-1^ *C*_*i*_ differentials at about ambient 300-350 μmol mol^-1^ CO_2_ in five amphistomatous species (Table 1). Considering the differential as the *C*_*i*_ gradient, they estimated the difference between calculated *C*_*i*_ and mean *C*_*i*_ to be 1/6 of the gradient or 3-10 μmol mol^-1^ for these amphistomatous species, according to the one-dimensional diffusion analysis. Their estimation depends on the location of calculated *C*_*i*_ which, in turn, depends on the diffusion path for water vapor because calculations assume the same pathway for CO_2_ and H_2_O in stomatal conductance. The diffusion path for stomatal conductance is then defined by the point where the gradients of water vapor starts (i.e. the conceptual evaporating surface), that is *W*_*i*_ (Eq. 2). Parkhurst et al. (1988) and Sharkey et al. (1982) considered this was right beneath stomata or stomatal cavity. However, this may not be true due to water vapor gradients and/or unsaturation of water vapor in the leaf airspace (see below). For the *C*_*i*_ differentials they observed, cuticle conductance might have an impact especially when stomatal conductance was small, yet they did not report *g*_*sw*_ (Table 1).

Dual sided open-flow data on amphistomatous leaves suggest that CO_2_ concentrations gradients are minimal across the leaf surface (Mott & O’Leary, 1984), however it is important to note that these data relied on the assumption that W_i_ is saturated at the substomatal cavity. Calculations for C_i_ are at the physical evaporating surfaces (W_i_), which is not necessarily in the substomatal cavity. Therefore, such data do not provide evidence against a CO_2_ concentration gradient *per se*, but rather that the CO_2_ concentration gradient is less than that required to cause a substantial difference between [CO_2_] at the evaporating surfaces on the adaxial and abaxial sides of the leaf.

As mentioned above, location of *W*_*i*_ is critical to define location of *C*_*i*_ through altering the diffusion path(-length) for stomatal conductance as illustrated in Fig. 2a. In general, *W*_*i*_ is considered to be saturated at *T*_*leaf*_ or 100% relative humidity (RH) throughout the airspace up until sub-stomatal cavity (shown as 100 in Fig. 2a). In this representation, calculated *C*_*i*_ is at the sub-stomatal cavity (*C*_*i,s*_), and the *C*_*i,s*_ is further reduced toward the mesophyll cell surface (*C*_*i,ias*_) due to finite *g*_*ias*_. When leaves are transpiring through stomata, evaporation essentially occurs on the cell surfaces exposed to the intercellular airspace (e.g., apoplast of the mesophyll cells). As for assimilation, a *W*_*i*_ gradient must exist from the evaporating surface to the stomatal cavity due to finite conductance to water vapor (shown as blue gradient on left hand side of Fig. 2a). In such case, the calculated *C*_*i*_ would be closer to the mesophyll cell surfaces where 100% RH occurs (i.e., *C*_*i,ias*_).

**Figure 2.**
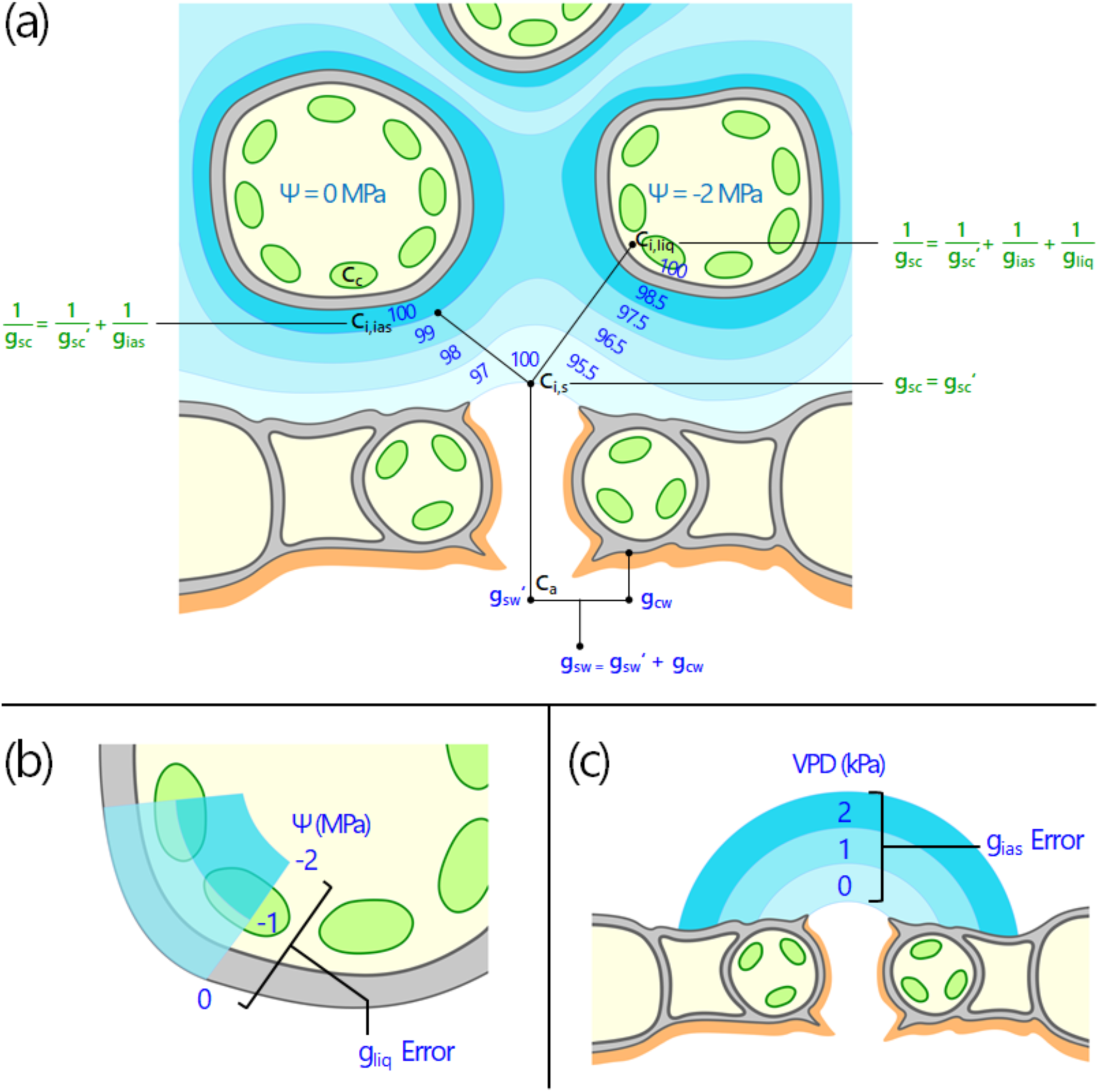
Implications of intercellular water vapor concentration (W_i_) as a percentage of saturation vapor pressure (VPD) for (a) conductance (g) calculations, and the effects of (b) water potential (Ψ) and (c) leaf to air vapor pressure deficit (VPD_leaf_). a) When Ψ = 0 MPa, W_i_ = 100% e_s_, and VPD = 0 kPa, then g represents stomatal conductance (g_s_). When Ψ = 0 MPa, W_i_ = 100% e_s_, and VPD > 0 kPa, then 1/g represents 1/g_s_ + 1/g_ias_ (intercellular airspace conductance). When Ψ < 0 MPa, W_i_ < 100% e_s_, and VPD > 0 kPa, then 1/g represents 1/g_s_ + 1/g_ias_ + 1/g_liq_ where g_liq_ is liquid conductance into the cell. Potential locations of C_i,es_ (CO_2_ concentration at the evaporating surface) are indicated by black lines. b) As Ψ decreases, the location of calculated W_i_ recedes further into the leaf and into the cell. c) As VPD_leaf_ increases, the location of W_i_ recedes away from the substomatal cavity. C_c_: chloroplastic CO_2_ concentration, C_ias_: CO_2_ concentration of the intercellular airspace.

#### Assumptions 3 – W_i_ saturation and Ψ_m,apo_ = 0 MPa

Besides *W*_*i*_ gradients, water potential of the water on the evaporating surface of the apoplast of the mesophyll cells (Ψ_m,apo_, the location most pertinent to C_i_) would affect the location of saturated *W*_*i*_ because RH over a solution is a function of Ψ of the solution as (Campbell & Norman, 1998): 

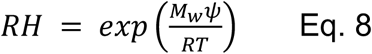

where *M*_*w*_ is the molecular weight of water (0.018 kg mol^-1^), Ψ is the water potential in J kg^-1^ (numerically equivalent to kPa), *R* is the universal gas constant (8.314 J K^-1^ mol^-1^), and *T* is the temperature in K. Eq. 8 indicates that RH in the airspace decreases as the Ψ_m,apo_ declines. If we assume that apoplastic and symplastic water potentials are in equilibrium in leaves, bulk water potential in leaf tissue (Ψ_leaf_), which we normally measure, may approximate the Ψ_apo_. We note that it is the water potential at the mesophyll apoplast that matters most for C_i_ calculations that are relevant for photosynthetically active tissues. Water potential of the bulk apoplast (Ψ_apo_) is composed of both mesophyll and bundle sheath apoplastic components (Ψ_m,apo_ and Ψ_b,apo_), and thus may be insufficient for calculating W_i_. However, the importance of this depends on the ratio of mesophyll to bundle sheath. If we assume that the bundle sheath + epidermal transpiration is small relative to mesophyll transpiration, then we can assume that Ψ_apo_ ≈ Ψ_m,apo_.

However if we assume for a moment that Ψ_leaf_ = Ψ_apo_, and Ψ_m,apo_ is the relevant value for calculating W_i_, for a leaf at night, Ψ_leaf_ can be as high as −0.1 MPa, corresponding to a *W*_*i*_ that is 99.9% RH, while a leaf at daytime with a Ψ_leaf_ of −2.0 MPa corresponds to a *W*_*i*_ of 98.5% RH (Eq. 8). Martínez-Vilalta et al. (2014) compiled a global dataset of water potential measurements, finding median predawn Ψ_leaf_ of −0.69 MPa (mean: −0.111 MPa, IQR: −0.13 – −0.34 MPa) and midday Ψ_leaf_ of −1.72 MPa (mean: −2.05 MPa, IQR: − 2.41 MPa – −1.23 MPa) corresponding to 99.5% (mean: 99.2%, IQR: 99.1 – 99.8 %) RH at predawn (assuming leaf temperatures of 25 °C) and 98.8% (mean: 98.5%, IQR: 98.3 – 99.1%) RH at midday. The effect of unsaturation on the calculation is illustrated on right hand side of Fig. 2a. 100% RH no longer exists in the intercellular airspace. Instead, it is located an imaginary point within the mesophyll cell (*C*_*i,liq*_), thereby causing the calculated *C*_*i*_ to be lower than the actual *C*_*i*_ in the airspace.

Some studies have suggested that the airspace could be unsaturated (Hygen, 1951, 1953; Slavik, 1958; Jarvis & Slatyer, 1970; Ward & Bunce, 1986; Egorov & Karpushkin, 1988; Karpushkin, 1994; Canny & Huang, 2006; Cernusak *et al*., 2018, 2019), while others have considered that effect of the Ψ_leaf_ is so small that the 100% RH can be assumed (Farquhar & Raschke, 1978; Jones & Higgs, 1980; Sharkey *et al*., 1982; Parkhurst et al., 1988; Buckley et al. 2017, Buckley and Sack 2019). However, Cernusak et al. (2018) recently estimated that the relative humidity could be as low as 77% and 87% (Ψ ≂ *–*35 MPa and *–*18 MPa, respectively) in *Pinus edulis* and *Juniperus monosperma* when the leaves opened stomata and actively photosynthesized. If true, unsaturation can have significant impact on the calculations at least in those species.

#### Assumption 4 – uniform stomatal apertures over leaf surfaces

Non-uniform stomatal apertures or stomatal ‘patchiness’ is another factor that could bias calculation of *C*_*i*_ (Downton et al., 1988; Terashima et al., 1988). They found patchy distribution of *A*_*net*_ throughout the leaves fed with ABA and proposed that if it is associated with stomatal patchiness, *C*_*i*_ would vary among patches and averaged *C*_*i*_ would be overestimated based on the conductance-weighed calculation (Eq. 5; Mansfield et al., 1990; Terashima, 1992; Buckley et al., 1997). Patchiness likely occurs in plants under water stresses that induce stomatal closure, however it is not a general phenomenon as it appears to depend on species, growth conditions, and how quickly the stress is imposed (Cheeseman 1991, Gimenez et al., 1992; Gunasekera & Berkowitz, 1992; Wise et al., 1992; Tezara et al., 1999; Mott & Buckley, 2000). A number of methods have been used to assess patchiness in conjunction with gas exchange measurements (Terashima, 1992): starch accumulation (Terashima et aI., 1988), autoradiography of fixed ^14^CO_2_ (Downton et aI., 1988a,b; Gunasekera & Berkowitz, 1992; Sharkey & Seemann, 1989; Wise et aI., 1992), fluorescence imaging (Daley et aI., 1989; Mott, 1995; Meyer & Genty, 1998). Although the results often have been attributed to patchy stomatal closure, these methods depend on photosynthetic metabolism and could as well reflect non-uniform metabolism (Lauer & Boyer, 1992; Wise et aI., 1992). Also, lateral CO_2_ diffusion and different stomatal behavior on both surfaces in amphistomatous leaves should affect the extent of observed patchiness. More independent and direct measurements of aperture/*g*_*sw*_ distributions, such as direct observations (Laisk et al., 1983; Van Gardingenet aI., 1989; Lawson et al., 1997) and thermal imaging (West et al., 2005; McAusland et al., 2013), may be preferable. The problems associated with stomatal patchiness are also attributable to *T*_*leaf*_ distributions that are difficult to measure accurately and are needed for *W*_*i*_. Direct *C*_*i*_ measurement can avoid these effects as it does not rely on conductance (Lauer & Boyer, 1992; Boyer & Kawamitsu, 2011).

#### Assumption 5 – no pressure gradients across leaf surfaces

Pressure gradients across the leaf surface would have direct impact on the concentration gradients because same atmospheric pressure is assumed for both inside and outside the leaf. There is evidence that humidity- and thermal-induced pressure gradients can exist across leaves, with data suggesting that this is the case in *Nelumbo* (Leuning, 1983; Dacey 1987) and *Nuphar lutea* (Dacey, 1981), however such pressure gradients may be associated most closely with aquatic plants (Steinberg, 1996). A modeling analysis suggests that intercellular airspace could be pressurized by up to 4 kPa—3.9% of 101.3 kPa for standard atmosphere—across the leaf (e.g. Steinberg, 1996), and that pressure gradients should increase with saturation of the intercellular airspace. If such pressurization can occur in terrestrial plants (which could happen under low g_ias_ values and increasing radiation loads as suggested by Steinberg, 1996), the possibility exists, therefore, that while W_i_ may not be equal to e_s_ *per se*, it may equal e_s_ at ambient air pressure if the leaf is pressurized and leaf RH < 100%, and or exceed the expected e_s_ if leaf RH is close to or greater than 100%. *W*_*i*_ calculated with the external ambient pressure would be lower than the actual *W*_*i*_ inside the leaf (i.e., the actual *W*_*i*_ is greater than 100% RH). To our knowledge, however, there are no studies demonstrating leaf to air pressure gradients in terrestrial plant species.

#### Assumption 6 – Fickian diffusion of CO_2_ and H_2_O

In the case of pressure gradients across the leaf in aquatic plants, the pressure gradients can be established because pore sizes are small enough that Knudsen diffusion is dominant over Fickian diffusion (Steinberg, 1996). Thus, it is possible that in some terrestrial species, the pore sizes could be sufficiently small as to cause Knudsen diffusion to occur, altering the diffusivity constants for CO_2_ and H_2_O, although stomatal pore size would need to be quite small for this effect (e.g. < 1 μm, Leuning, 1983), and the ratio of Knudsen diffusion coefficients for H_2_O and CO_2_ would be 1.56 as the ratios are dependent on pore size and molecular mass. Thus, while Knudsen diffusion may occur in some cases, the assumption of Fickian diffusion is likely sufficient for terrestrial plants in most cases.

#### Assumption 7 – one dimension approximates the three-dimensional leaf

The equations used in gas exchange are typically one dimensional, and it is generally assumed that this is sufficient to capture the behaviour of the leaf area measured through gas exchange. However, this may be insufficient (Parkhurst, 1977) and three-dimensional models predict that some gas exchange traits could be strongly affected (Parkhurst, 1994; Earles et al., 2018). Furthermore, three dimensional models predict mechanisms behind some of the responses observed in mesophyll conductance to CO_2_ (*g*_*m*_) (Tholen & Zhu, 2011).

### Implications of broken assumptions – where is C_i_?

Most of the assumptions listed above essentially relate to meaning of stomatal conductance—source of transpiration, diffusion path, behavior and diffusive capacity of stomata. While cuticular CO_2_ movement is very small and probably negligible in *A*_*net*_ considerable cuticular water loss occurs in transpiration, and so *g*_*sw*_ should include *g*_*cw*_. Because stomatal and cuticular transpiration occur in parallel cuticle conductance is additive to stomatal conductance as (Fig. 2a): 

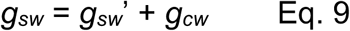

where *g*_*sw*_ is what we calculate according to Eq. 2 where *E* includes stomatal and cuticular transpiration whereas *g*_*sw*_’ accounts for only stomatal transpiration. Eq. 9 shows that the *g*_*sw*_ is overestimated by *g*_*cw*_ (Fig. 2a). Also, the effect of *g*_*cw*_ on the calculation is expected to be greater with the larger *g*_*cw*_ and smaller *g*_*sw*_*’*, both of which increase the proportion of *g*_*cw*_ to *g*_*sw*_.

Parkhurst (1994) and co-workers suggested C_i_ as we calculate it is better represented by C_i,s_ for C_i_ at the stomatal cavity based on the calculations used (Fig. 2a). Parkhurst (1994) predicted that the degree of C_i_ over-estimation relative to the average *C*_*i*_ would be greater for hypostomatous leaves rather than amphistomatous leaves, and further argues that even dual-sided gas exchange measurements can only measure C_i,s_, the average *C*_*i*_. However, calculated *C*_*i*_ could be deeper than they assumed. Diffusion path of *g*_*sw*_ affects location of *C*_*i*_ and is potentially complicated because where a *W*_*i*_ gradient occurs and where *W*_*i*_ is saturated (100% RH) might change with microenvironment and water status in leaves. To disentangle these effects, it is helpful to find 100% RH within the leaf because it defines the starting point of *g*_*sw*_ as well as the end point of *g*_*sc*_ which sets the location of *C*_*i*_. In Fig. 2a, stomatal conductance to CO_2_ (*g*_*sc*_) for each *C*_*i*_ site as well as *W*_*i*_ are indicated.

When there is no *W*_*i*_ gradient and the Ψ_m,apo_ is zero, sub-stomatal cavity may be saturated (center of Fig. 2a). In this scenario, *C*_*i*_ would be calculated as *C*_*i,s*_ with the *g*_*sc*_ accounting only stomatal path (*g*_*sc*_ = *g*_*sc*_’). We note that the substomatal cavity may not be saturated if the cuticle extends into the substomatal cavity as it does for *Tradescantia virginiana* (Nonami et al. 1991). When the *W*_*i*_ gradients exist and the Ψ is zero, 100% RH may be found at the mesophyll cell surface (left hand side of Fig. 2a). Now, the calculated *C*_*i*_ would indicate the *C*_*i,ias*_ with the *g*_*sc*_ accounting for the stomatal plus intercellular pathway from the stomatal cavity to the mesophyll cell surface 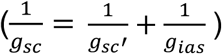. When the Ψ_m,apo_ is negative (right hand side of Fig. 2a), 100% RH may be within the mesophyll cell. In this case, the *g*_*sc*_ partially includes CO_2_ diffusion path in the liquid phase in addition to the air phase 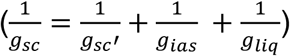, and the *C*_*i*_ would be calculated as *C*_*i,liq*_. The *C*_*i,liq*_ would be located in the liquid path from the cell wall surface (*C*_*i,ias*_) to chloroplast stroma (*C*_*c*_) depending on where the assumed *W*_*i*_ is located. Importantly, *g*_*ias*_ and *g*_*liq*_ usually resides in the mesophyll conductance (*g*_*m*_) as (Evans et al., 2009): 

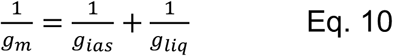

where *g*_*m*_ is defined as: 

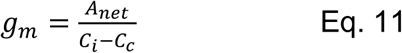

Clearly, *g*_*m*_ is affected by *g*_*cw*_ and *W*_*i*_ which break the assumptions of *g*_*sc*_ = *g*_*sc*_*’* (i.e., *C*_*i*_ = *C*_*i,s*_, the C_i_ in the substomatal cavity). If *g*_*ias*_ is included in *g*_*sc*_ when saturated *W*_*i*_ occurs at the mesophyll cell surface, calculated *C*_*i*_ would be *C*_*i,ias*_ rather than *C*_*i,s*_ and *g*_*m*_ might be calculated to be strictly liquid-phase conductance (*g*_*m*_ = *g*_*liq*_). Furthermore, some portion of *g*_*liq*_ is mis-assigned to *g*_*sc*_ as the Ψ_apo_ ‘pulls’ the *C*_*i,liq*_ deeper into the mesophyll cells (*g*_*liq*_ error in Fig. 2b). Consequently, decreasing the path-length for CO_2_ overestimates the apparent *g*_*m*_ (Eq. 10. Even if sub-stomatal *W*_*i*_ is saturated, the vapor pressure difference between air and leaf (VPD_leaf_) may ‘push’ the *W*_*i*_ deeper into the airspace by making the *W*_*i*_ gradients steeper (Fig. 2c). Then, *g*_*m*_ would also be overestimated by reducing some portion of path-length in the airspace (*g*_*ias*_ error in Fig. 2c). These ‘pushing’ and ‘pulling’ effect can happen either independently or simultaneously or in coordinated manner when environmental water demand is excessive (e.g., under drought). We can see by this illustration that only under very specific circumstances can our current assumptions about *W*_*i*_ provide us with ‘true’ *g*_*s*_, *C*_*i*_, and *g*_*m*_.

In the following sections, we model the implications when some of these assumptions fail, reinterpret data from the literature in several case studies on mesophyll conductance and the photosynthetic CO_2_ response, and propagate different rates of cuticular water loss, *W*_*i*_ gradients, and unsaturation through the data.

#### Modeling the implications of cuticle conductance

##### Re-calculations of C_i_, intrinsic water use efficiency, mesophyll conductance

Analogous to Eq. 5, *C*_*i*_ was recalculated (*C*_*i,cuticle*_) with the actual *g*_*sw*_’ in Eq. 9 as: 

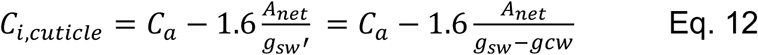

For *C*_*a*_, we assumed infinite boundary layer conductance as most papers do not report enough information to recalculate or extract boundary layer conductance and gas exchange chambers are designed to minimize boundary layers. In Eq. 12, we also assumed that cuticle conductance for CO_2_ was negligible (*g*_*cc*_ = 0) because the effect of the *g*_*cc*_ on the *A*_*net*_ has been often undetectably small under experimental *C*_*a*_ levels (Boyer et al., 1997; Tominaga et al., 2018).

To propagate *g*_*cw*_ through the modeling, we fixed *g*_*cw*_ as a proportion of stomatal conductance (0, 1 x 10^−5^, 1 x 10^−4^, 1 x 10^−3^, 1 x 10^−2^, 5 x 10^−2^, 1 x 10^−1^, and 2.5 x 10^−1^) at the lowest light intensity for light response curves (see below), 25 °C for temperature response curves (see below), and 400 μmol mol^-1^ CO_2_ for the CO_2_ response curves (see below).

Introducing *g*_*cw*_ into a gas exchange approach to plant water balance has implications for how we define water use efficiency, as we can partition out stomatal and cuticular water use efficiencies. By separating out cuticle and stomatal water loss components, we can better understand the immediate cause as to why plants vary in water use efficiency (i.e. stomatal versus cuticular components). This partitioning could then be used to inform crop breeding for further enhancing water use efficiency. We recalculated intrinsic water use efficiency (*iWUE*) as: 

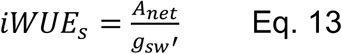

where *iWUE*_*s*_ is intrinsic stomatal water use efficiency, (μmol CO_2_ mol^-1^ H_2_O). To calculate the effects of including cuticular conductance on water use efficiency, we used a representative steady-state *A*-*C*_*i*_ curve for *Populus deltoides* from Stinziano *et al*. (2017) and we propagated *g*_*cw*_ as a proportion of *g*_*sw*_ at a reference [CO_2_] of 400 μmol mol^-1^. For this propagation, we recalculated *g*_*sc*_ according to the standard procedure (Li-Cor, 2019): 

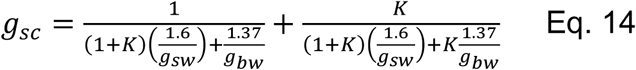

where *K* is the ratio of stomata on the adaxial to the abaxial surface of the leaf (assumed to be equal to 1), and g_bw_ is the boundary layer conductance to water. Accounting for cuticular conductance in the calculations of water use efficiency leads to an increase of up to 20% in iWUE when cuticular conductance is high (Fig. 3).

**Figure 3.**
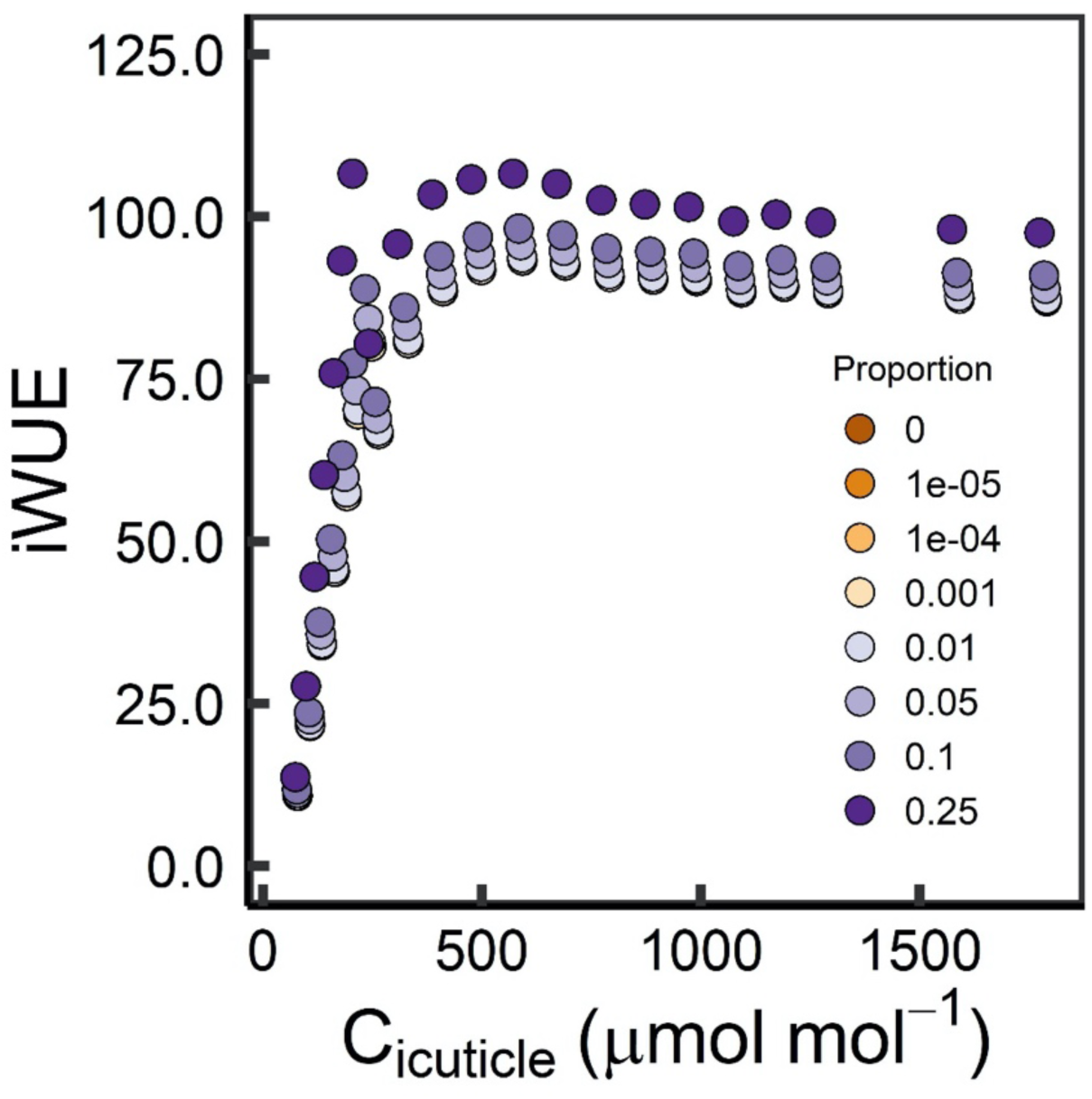
Water use efficiency as a function of intercellular [CO_2_] accounting for cuticular conductance (C_icuticle_), and the relationship with the proportion of stomatal conductance attributed to cuticular conductance. iWUE: intrinsic water use efficiency.

#### Implications of cuticular conductance on the interpretation of mesophyll conductance data

##### g_m_ Calculations

Since C_c_ is also dependent on C_i_, we need to set out *a priori* predictions of how changes in C_i_ would affect C_c_. To predict this effect, we started with the equation describing the online isotope discrimination from (Farquhar *et al*., 1982) as modified by (Wingate *et al*., 2007): 

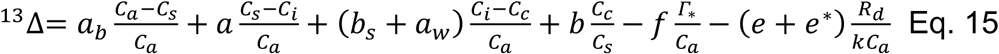

where ^13^Δ is the predicted net ^13^C discrimination, a_b_ is the ^13^C fractionation due to diffusion through the boundary layer, a is the ^13^C fractionation due to diffusion through the stomata, b_s_ is the ^13^C fractionation due to CO_2_ solubilization, a_w_ is the ^13^C fractionation during diffusion in water, b is net ^13^C fractionation during carboxylation by rubisco and PEP carboxylase, f is the ^13^C fractionation due to photorespiration, e is the ^13^C fractionation due to decarboxylation, e^*^ is apparent ^13^C discrimination during decarboxylation, k is carboxylation efficiency, G_*_ is the photorespiratory CO_2_ compensation point, and R_d_ is the rate of respiration in the dark. Note that for the sake of simplicity, we ignore all ternary interactions here (Farquhar and Cernusak, 2012) as it becomes an unnecessarily complex for demonstrating the reliance of g_m_ on C_i_ for this review (see below). We can rearrange this for C_c_ to obtain: 

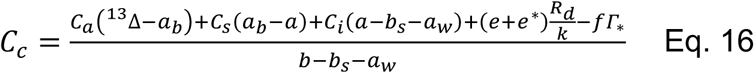

Now suppose we have C_i,standard_ and C_i,cuticle_, and want to calculate the difference between C_c,standard_ (C_c_ determined without g_c_) and C_c,cuticle_. By calculating the difference, most terms in the above Eq. cancel out (even the term with k, which should be the same in theory; see Appendix A for details), leaving us with: 

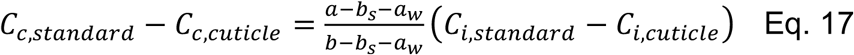

Which can be further rearranged to: 

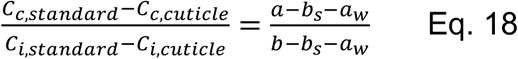

In this way, the difference in C_c_ can be calculated using a ratio of fractionation constants and the difference in C_i_. Since a is typically assumed to be 4.4 ‰, b_s_ is assumed to be 1.1 ‰ at 25 °C (Vogel, 1980), a_w_ is assumed to be 0.7 ‰, and b is assumed to be between 27 and 30 ‰, then the difference in C_c_ values should be between 9.2 and 10.3 % of the difference in C_i_ values at 25 °C. For the g_m_ calculations, we took a conservative approach and assumed that the difference in C_c_ was 9.2% of the difference in C_i_ values.

Including the ternary effects from Farquhar and Cernusak (2012) makes *a priori* predictions of the effect of cuticle conductance on C_c_ more difficult. Describing the discrimination: 

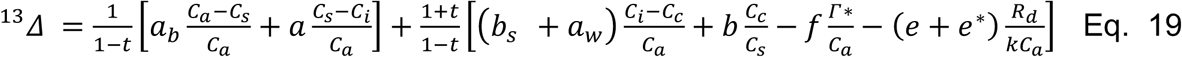

Where t is the ternary term described by: 

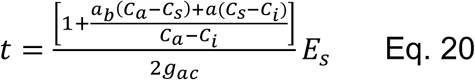

Where g_ac_ is the total conductance to CO_2_ diffusion. Note that C_i_, E_s_, and g_ac_ all need to be corrected for g_sw_ and g_c_ occurring in parallel. Since g_c_ affects nearly every component of t, calculations using ^13^C discrimination may be highly sensitive to g_c_ when considering ternary interactions. However, if we assume a C_a_ of 400 μmol mol^-1^, g_bw_ of 2 mol m^-2^ s^-1^, g_sw_ of 0.15 mol m^-2^ s^-1^, leaf-to-air vapor pressure deficit of 1.0 kPa, A_net_ of 15 μmol m^-2^ s^-1^, t changes value from 2.6547 in the case of no cuticle conductance up to 2.659 in the case where 10% of g_sw_ is attributed to cuticle conductance. Thus, the ternary calculations may be minimally sensitive to cuticle conductance. We can rearrange Eq. 20 for C_c_: 

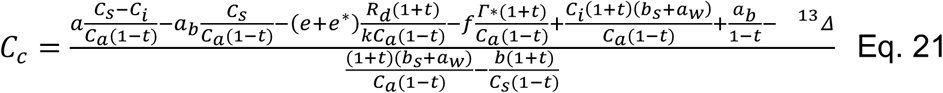

We can see that the effect of cuticle conductance affects nearly every term in the equations. And solving for the difference between C_c_ without cuticle conductance (C_c,s_) and with cuticle conductance (C_c,c_) we get (See Appendix B for derivation): 

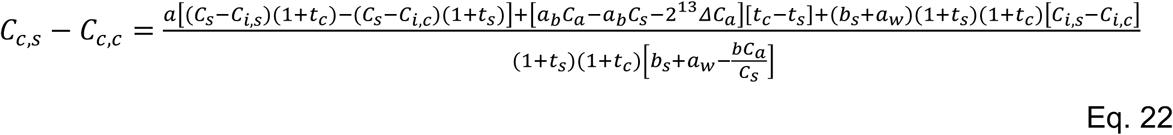

where C_i,s_ is C_i_ without cuticle conductance, C_i,c_ is C_i_ accounting for cuticle conductance, t_s_ is the ternary equation without cuticle conductance, and t_c_ is the ternary equation with cuticle conductance. For the purposes of this review, however, we have not included the ternary effects into our modeling.

#### Reinterpreting gm data

We reinterpreted g_m_ by propagating a g_cw_ into the g_sw_ data through recalculating *C*_*i*_ from Eq. 12 and gm from Eq. 11. We fixed *g*_*cw*_ as a proportion of stomatal conductance (0, 1 x 10^−5^, 1 x 10^−4^, 1 x 10^−3^, 1 x 10^−2^, 5 x 10^−2^, 1 x 10^−1^, and 2.5 x 10^−1^) at the lowest light intensity for light response curves (see below), 25 °C for temperature response curves (see below), and 400 μmol mol^-1^ CO_2_ for the CO_2_ response curves (see below). For reinterpreting the g_m_ temperature response data from Scafaro et al. (2011), we included temperature response functions of g_cw_ obtained from Riederer and Schreiber (2001) fitting an Arrhenius equation on either side of the breakpoint in the Arrhenius plot: 

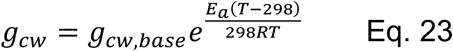

where g_cw,base_ was determined as above, the highest and lowest E_a_ (25,215 J mol^-1^ with a breakpoint at 35 °C to 85,171 J mol^-1^; 20,145 J mol^-1^ with a breakpoint at 30 °C to 69,856 J mol^-1^, respectively) from Riederer and Schreiber (2001) were used, T is the leaf temperature in K, and R is the universal gas constant (8.314 J mol^-1^ K^-1^). For g_m_ light response data, we calculated g_cw_ based on the g_a_ data from the lowest light intensity to minimize issues with g_cw_ exceeding g_a_. In nearly all cases, g_cw_ substantially affected g_m_, with notable changes occurring when g_c_ exceeds ∼1% of g_s_ (Fig. 4b, d, f). In regard to environmental responses of g_m_, light and temperature responses are much more sensitive to g_cw_ than CO_2_ response (Fig. 4b, d, f). It is important to note that g_m_ appears most sensitive to g_cw_ for C_i_ < 500 μmol mol^-1^.

**Figure 4.**
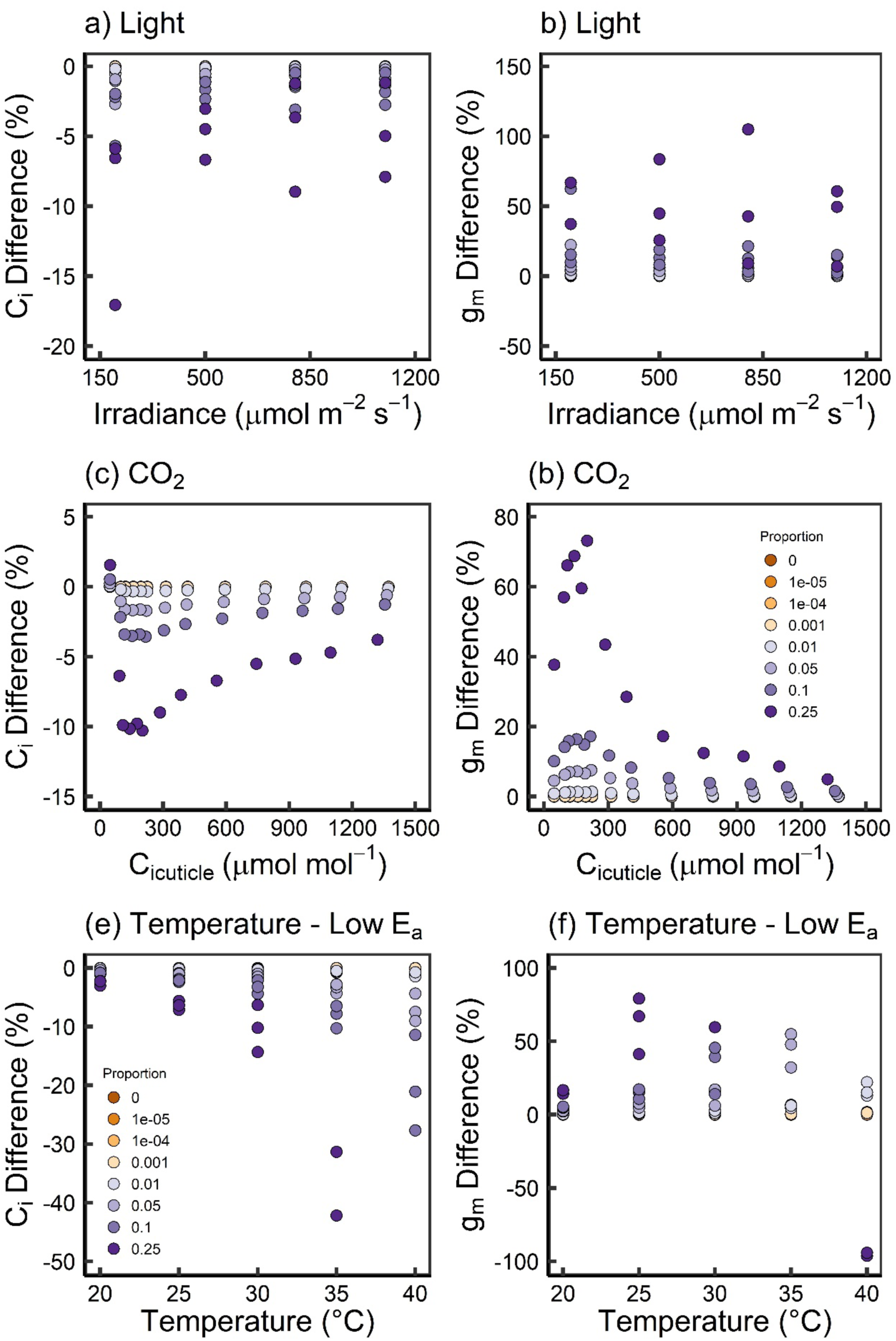
The sensitivity of intercellular [CO_2_] (C_i_) (a, c, e) and mesophyll conductance (g_m_) (b, d, f) to the proportion of stomatal conductance attributed to cuticle conductance. (a) Data from Douthe et al. (2011); (b) Data from Vrabl et al. (2009); (c, d) Data from Scafaro et al. (2011) assuming (c) the highest temperature sensitivity of cuticular conductance or (d) the lowest temperature sensitivity of cuticular conductance from Riederer and Schreiber (2001). For unrestricted axes and C_i_ comparisons, see Figs. S1, S2; for g_m_ comparisons, see Figs. S3, S4, S5).

#### Reinterpreting A-C_i_ data

We used a representative steady-state A-C_i_ response from *Populus deltoides* (Stinziano *et al*., 2017). A-C_i_ data were recalculated by assuming: 1) constant g_cw_ across all C_i_, and 2) g_cw_ was 0 to 25% (in increments of 0.1%) of g_sw_ at C_a_ of 400 μmol mol^-1^. All A-C_i_ curves were fit using the ‘bilinear’ approach (which treats the FvCB model as a change-point model during curve fitting similar to Gu et al., 2010) of the {plantecophys} package in R (Duursma, 2015). For species that had temperature response curve measured, we fit the modified Arrhenius model to the maximum Rubisco carboxylation (V_cmax_) and maximum electron transport (J_max_) rates.

Accounting for cuticle conductance causes a decrease in the calculated value of C_i_, such that current calculations methods are systematically overestimating C_i_ (Fig. 4a, c, e). These differences are least pronounced for A-C_i_ curve data (Fig. 4c), and most pronounced for temperature response data. In some instances, the C_i_ calculations breakdown when g_cw_ approaches the value of g_sw_. Accounting for g_cw_ alters the shape of the A-C_i_ response, and increases fitted values for V_cmax_ and J_max_, although differences appear negligible until g_cw_ is ∼1% of g_sw_ (Fig. 5). In light of the interpretations of Parkhurst (1994), we modelled the impact of combined CO_2_ gradients and cuticle conductance on the perceived A/C_i_ response.

**Figure 5.**
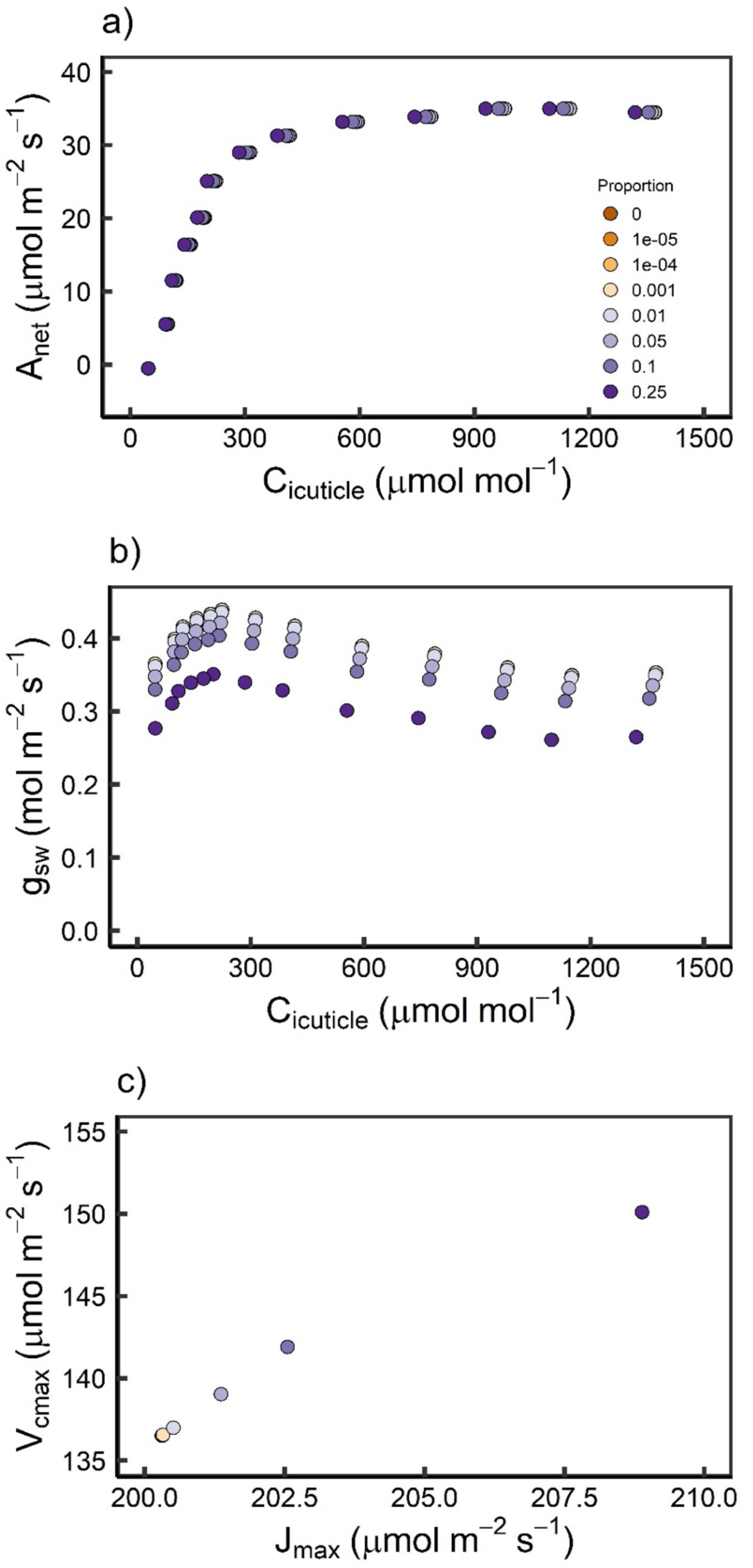
a) Response of A_net_ to C_i_ under different g_cw_ scenarios. b) Response of g_sw_ to C_i_ under different g_cw_ scenarios. Solid lines indicate value of g_cw_ across curve. c) Response of V_cmax_ to J_max_ as a function of the proportion of g_sw_ attributed to g_cw_. Re-interpreted A-C_i_ data extracted from Vrabl et al. (2009). A_net_: net CO_2_ assimilation; C_i_: intercellular CO_2_ concentration; g_cw_: cuticle conductance to water; g_sw_: stomatal conductance to water; J_max_: maximum rate of electron transport to RuBP regeneration; Proportion: proportion of g_sw_ attributed to g_cw_ at a reference CO_2_ of 400 μmol mol^-1^; V_cmax_: maximum rate of rubisco carboxylation capacity.

#### Modeling g_ias_ and W_i_ effects on gas exchange parameters

Considering the Eq. describing the surface to intercellular CO_2_ concentration gradient: 

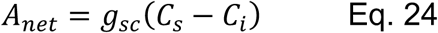

where g_sc_ is stomatal conductance to CO_2_, C_s_ is the CO_2_ concentration at the leaf surface, and C_i_ is intercellular airspace CO_2_ concentration. Eq. 24 can misrepresent the process, which according to Parkhurst (1994) would be: 

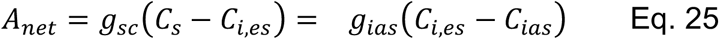

where C_i,es_ is the CO_2_ concentration at the site of evaporating surfaces, g_ias_ is the conductance of CO_2_ from the evaporative surfaces to the intercellular airspace, and C_ias_ is the concentration of CO_2_ in the intercellular airspace. Note that under the Parkhurst (1994) definition, C_ias_ could represent a point anywhere in the intercellular airspace, while C_i,es_ is assumed to be closer to the substomatal cavity than C_ias_. If the evaporating surface is at the mesophyll cell surface, then Eq. 25 would have to account for this reversed order and be re-written as: 

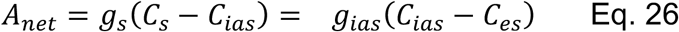

Furthermore, g_s_ needs to be corrected for cuticular conductance. Therefore, we can describe the conductance of CO_2_ and water from the leaf surface to the intercellular airspace (g_s,ias_) according to: 

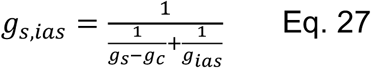

We can further include mesophyll conductance, g_m_, to calculate total conductance of CO_2_ and water from the leaf surface to the chloroplast (g_t_): 

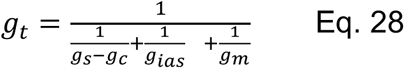

We can then model the implications of g_c_ and g_ias_ on C_i_ by varying their values. We linked Eq. 28 to a leaf-level model of photosynthesis: 

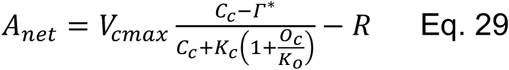

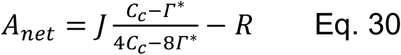

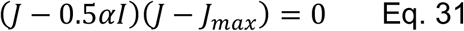

where V_cmax_ is maximum rate of Rubisco carboxylation, O_c_ is the oxygen concentration in the chloroplast (210 mmol mol^-1^), K_c_ is the Michaelis-Menten constant for Rubisco carboxylation, K_o_ is the Michaelis-Menten constant for Rubisco oxygenation, J_max_ is the maximum rate of electron transport, J is the rate of electron transport, † is the proportion of irradiance (I) absorbed by the leaf, R is respiration, and G* is the photorespiratory CO_2_ compensation point. All values (except for V_cmax_ and J_max_) were obtained from Bernacchi *et al*. (2001).

We assumed a V_cmax_ and J_max_ of 100 and 200, respectively, and modelled under light saturating conditions such that J = J_max_. We modelled from a C_s_ of 50 to 2000 in 50 ppm intervals. For calculating E_s_ (to represent the ‘measured’ transpiration from a gas exchange cuvette) we used the following equation: 

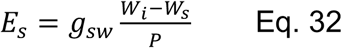

where E_s_ is stomatal transpiration, g_sw_ is stomatal conductance (ranging from 0.03 to 2.00 mol m^-2^ s^-1^), W_i_ is the water concentration inside the leaf (assumed to be 100% saturation vapor pressure for this initial calculation, which was calculated according to Cernusak et al., 2018), W_s_ is the water vapor concentration at the leaf surface (assumed to be 50% saturation vapor pressure), and P is atmospheric pressure (assumed to be 100 kPa). We assumed leaf temperature was equal to air temperature of 298 K. Once E_s_ was calculated, we then altered our assumptions about W_i_, changing it to 99% and 90% of saturation vapor pressure. Then for each different W_i_ scenario, we set g_cw_ to either 0 or 0.01, and g_ias_ to either 1.00 (Mott, 1988) or infinity. The C_i_ obtained when W_i_ = 100% saturation vapor pressure, g_cw_ = 0 and g_ias_ = infinity was used as the reference C_i_. A_net_ was then modelled to obtain A/C_i_ responses (using both reference C_i_ and the C_i_ obtained from each combination of W_i_, g_cw_ and g_ias_) which were then fit using {plantecophys} (Duursma, 2015) in R (R Core Team, 2018) to obtain V_cmax_ estimates.

Modeling the effects of g_cw_ and g_ias_ across a range of reference g_sw_ (i.e. the g_s_ ‘measured’ using a typical open-flow gas exchange system), we see that g_c_ has the greatest impact at low g_sw_, with a negligible effect when g_cw_/g_sw_ < 1% at a C_s_ of 400 ppm (Fig. 6). Finite g_ias_, however, has a much larger impact on C_i_, with its effect size increasing with g_sw_ (Fig. 6). This explains the Ci discrepancies observed in Table 1— larger discrepancy with low g_sw_ and smaller discrepancy with high g_sw_. It is also possible that the discrepancies relate to C_ias_ being directly measured deeper in the intercellular airspace than the location of the evaporating surface such that the calculated C_i_ is C_i,es_ and the differences are due to how the quantities are defined. In the case of leaves treated with ABA (e.g. Boyer 2015a,b; Tominaga & Kawamitsu, 2015a) or stress induced stomatal closure, g_cw_ could account for the majority of the impact, since the limit of the CO_2_ gradient-related deviation in calculated and real C_i_ tends towards 0 as measured g_sw_ approaches 0. Looking at W_i_, the impact of W_i_ assumptions is evident. A 1% reduction in W_i_ increases the C_i_ discrepancy by a few ppm (Fig. 6b), and a 10% reduction causes changes the discrepancy by over 20 ppm in some cases (Fig. 6c).

**Figure 6.**
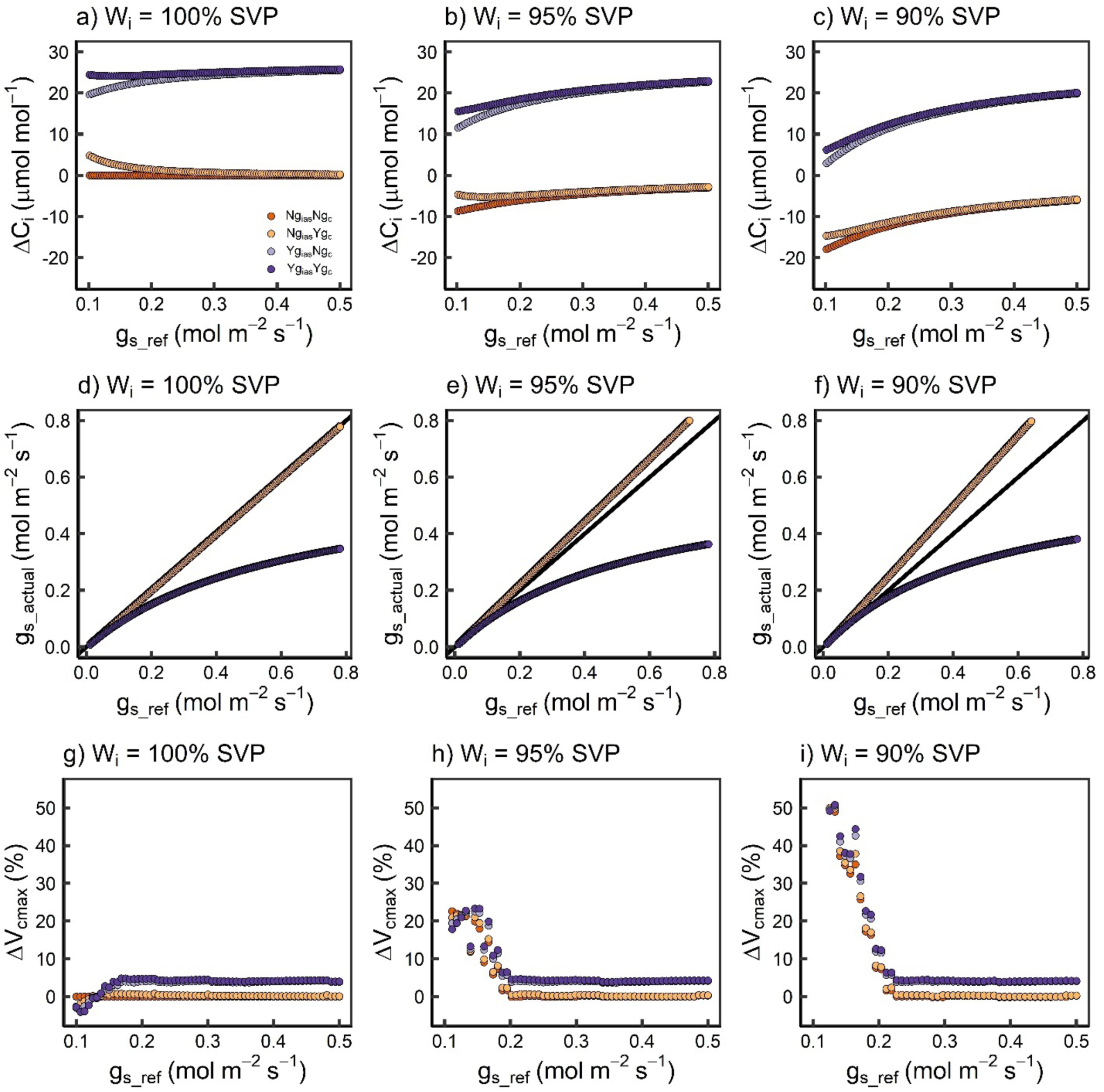
Effects of CO_2_ gradients (finite g_ias_) and cuticle conductance (g_c_) on calculated C_i_ (ΔC_i_) (a, b, c), g_s_ (d, e, f), and ΔV_cmax_ (g, h, i), when intercellular water vapor concentration (W_i_) is at 100% (a, d, g), 95% (b, e, h), and 90% (c, f, i) of saturation vapor pressure (e_s_). g_s_actual_: actual stomatal conductance given the assumptions; g_s_ref_: reference g_s_ where g_c_ = 0, g_i_ mmol m^-2^ s^-1^_s_ = infinity, and W_i_ = 100% e_s_; Ng_ias_: no CO_2_ gradient; Yg_ias_: CO_2_ gradient present, g_ias_ = 1000 mmol m^-2^ s^-1^; Ng_c_: g_c_ = 0 mmol m^-2^ s^-1^; Yg_c_: g_c_ = 10 mmol m^-2^ s^-1^; ΔV_cmax_: percent change in maximum rate of Rubisco carboxylation.

If we fit A/C_i_ curves in the presence of g_cw_ and finite g_ias_ and use the Moss & Rawlins (1963) assumptions, g_c_ has a relatively small impact on V_cmax_, but is important in cases where g_cw_ > 5% of g_s_, while g_ias_ causes a large depression in V_cmax_ across all g_sw_ used in simulations (Fig. 6g, h, i). As vapor pressure in the leaf is reduced from 100%, V_cmax_ estimates increase (Fig. 6g, h, i). Interestingly, with g_cw_ > 0, g_ias_ < ∞, internal vapor pressure < saturation vapor pressure and high g_sw_, V_cmax_ estimations are close to the value used in the model (Fig. 6i). Based on these modeling analyses, the impact of finite g_ias_ may be of greater concern when estimating gas exchange parameters than g_c_, and many of the large g_c_ values reported using C_i_ differentials between calculated and measured values may in fact be partially attributed to a finite g_ias_. It is crucial to note, however, that cuticle water fluxes have been reported up to 65% of total water flux across a leaf (Šantrůček et al., 2004), and the relative influence of g_ias_ and g_cw_ depend on the relative value of g_sw_. Given our modeling results showing the different impact of g_cw_ and g_ias_ on gas exchange data, it may be possible to construct a model capable of estimating g_cw_ and g_ias_ from a data set. This would allow proper attribution of C_i_ differentials to g_cw_ versus g_ias_.

Given the impact when all three assumptions test above are violated, it is possible that many (or even most) estimates of apparent V_cmax_ in the literature may still be ‘correct’ for the wrong reasons. However, we would like to note important assumptions made in our modeling: 1) resistances within the leaf are additive (which may not hold; Parkhurst, 1984), 2) Fickian (rather than Knudsen, which may occur; Dacey, 1987) diffusion governs gas diffusion from outside to inside the leaf, 3) the leaf is treated one-dimensionally rather than three-dimensionally (which will affect calculations: Parkhurst, 1977; Earles et al., 2018), 4) the air pressure differential from outside to inside the leaf is 0 (evidence suggested this may not be correct, at least in the extreme case of lotus, *Nelumbo*; Leuning, 1983; Dacey, 1987).

### Solving the failed assumption

#### When does it (not) work?

Cuticular conductance has largely been assumed negligible and is often ignored in gas exchange measurements, while CO_2_ gradients are largely ignored - however this may be due to partitioning g_ias_ into g_m_ (Evans et al. 1994), reducing the need to consider g_ias_ when C_i_ is taken as C_i,es_. To date, many of the Moss and Rawlins (1963) assumptions have been shown to be incorrect in at least some cases (e.g. Hygen, 1951, 1953; Slavik, 1958; Jarvis & Slatyer, 1970; Leuning, 1983; Parkhurst, 1984; Ward & Bunce, 1986; Dacey, 1987; Egorov & Karpushkin, 1988; Long et al., 1989; Karpushkin, 1994; Boyer et al., 1997; Meyer and Genty, 1998; Šantrůček et al., 2004; Canny & Huang, 2006; Boyer 2015a; Boyer, 2015b; Tominaga and Kawamitsu, 2015a; Cernusak et al., 2018; Tominaga et al., 2018, Cernusak et al. 2019). We summarize the assumptions and their expected impact on C_i_ calculations in Fig. 8. However, it is important to note that the assumptions have allowed major breakthroughs in our understanding of plant physiology.

Our analysis of the effects of g_cw_ on gas exchange measurements suggests that C_i_ is relatively unaffected when g_cw_ is less than 1% of g_sw_ across a range of irradiance, [CO_2_], and temperature, and has a relatively minor effect on fitted values of V_cmax_ and J_max_. Given that most values of g_cw_ measured to date are relatively low, and assuming g_cw_ was measured correctly, it is likely below the 1% threshold in unstressed plants, especially crops (Table 2; Schuster *et al*., 2017), this would explain why the Moss and Rawlins (1963) assumption that g_cw_ = 0 has been successful in advancing our understanding of photosynthesis over the past six decades. In regard to g_m_, accounting for g_cw_ increases the value of g_m_, however such effects are small across irradiance and [CO_2_] when g_cw_ is at or below 1% g_sw_ but become particularly important for the temperature response of mesophyll conductance. When g_cw_ exceeds 1% of g_sw_, the calculations for mesophyll conductance broke down for the modeling, giving extremely high and/or negative values for g_m_, which is related to C_i,cuticle_ dropping close to or below C_c_. Pons et al. (2009) recommended accounting for g_cw_ in g_m_ measurements, and our modeling suggests that this is critical when looking at the temperature response of g_m_, and in cases where g_cw_ is very high relative to g_sw_. We may thus expect significant errors in gas exchange calculations when g_sw_ is low (i.e. low light, drought, and high VPD conditions), and/or when g_cw_ is high (i.e. high temperature, well-watered plants). Furthermore, considering chlorophyll fluorescence-based estimates of g_m_ (i.e. Harley et al., 1992), sensitivity of g_m_ should be similar to the isotopic method as it is calculated via Eq. 23, with the added caveat that C_c_ becomes sensitive to the estimate of the photorespiratory CO_2_ compensation point (G*). Since G* can be estimated from gas exchange or Rubisco kinetics, the sensitivity of calculated g_m_ to g_cw_ via the variable J method will depend on how G* is measured. In this regard, interpreting data from drought and temperature stress experiments should proceed with caution if g_cw_ is ignored.

CO_2_ gradients within leaves have ontological consequences for gas exchange measurements, in particular, the meaning of C_i_ (Parkhurst, 1994). If typical C_i_ estimates are taken to be C_i,es_ measurements, then the implications of a finite g_ias_ on data derived from the C_i_ estimates are minimal, since g_ias_ is often subsumed into g_m_ (Evans et al., 1994). If C_i,es_ occurs at the surface of the mesophyll cells, then g_ias_ will have no impact on C_i_ estimates since the g_sw_ calculation occurs at the location of the evaporating surface (Parkhurst, 1994). However, if the C_i,es_ is located closer to the stomata than C_ias_ (i.e. if the evaporating surface is not the mesophyll cell surface), C_ias_ would then be located closer to the mesophyll cells than C_i,es_. Our modeling of finite g_ias_ represents this case and demonstrates that the implications of g_ias_ on data derived from such C_i_ estimates can be quite large, causing C_i_ estimates to differ by >10 μmol mol^-1^, and g_sw_ to be reduced by more than 50% (Fig. 6). To reiterate, subsuming g_ias_ into g_m_ eliminates the consequences of g_ias_ for ‘C_i_’ estimates under conditions where C_ias_ lies closer to the mesophyll cells than C_i,es_. If C_i,es_ is at the mesophyll cell surface, then g_ias_ is already accounted for in g_sw_ calculations. As long as it is recognized that C_i_ estimates represent C_i,es_ (Parkhurst, 1994) and that all derived parameters are apparent parameters on a C_i,es_-basis, g_ias_ poses minimal issues to the interpretation of C_i_ data. This becomes an issue however, in cases where the location of the evaporating surface differs between species or treatment groups (e.g. control versus drought stress). The same would be true for parameters derived from C_ias_ estimates if the rest of g_m_ were ignored. Note that this underscores the importance of knowing the location of the C_i_ calculation. However, it appears that g_ias_ has minimal consequences for g_m_ relative to g_cw_ at low values of g_sw_ (Fig. 7), with g_ias_ shifting the g_sw_ value at which g_cw_ has the greatest impact on g_m_. Reducing W_i_ tends to reduce the impact of g_cw_ and g_ias_ on g_m_ (Fig. 7). We also calculated the theoretical maximum values for g_ias_ based on leaf thicknesses from Onoda et al. (2011) to estimate diffusion distances, along with biophysical equations to calculate conductance (Massman, 1998; Campbell & Norman, 1998) (see Supplementary Methods for more information on the calculations). We calculated that the median maximum theoretical values of g_ias_ for an amphistomatous and hypostomatous leaf is 24 and 3 mol m^-2^ s^-1^, respectively (Fig. 1c).

**Figure 7.**
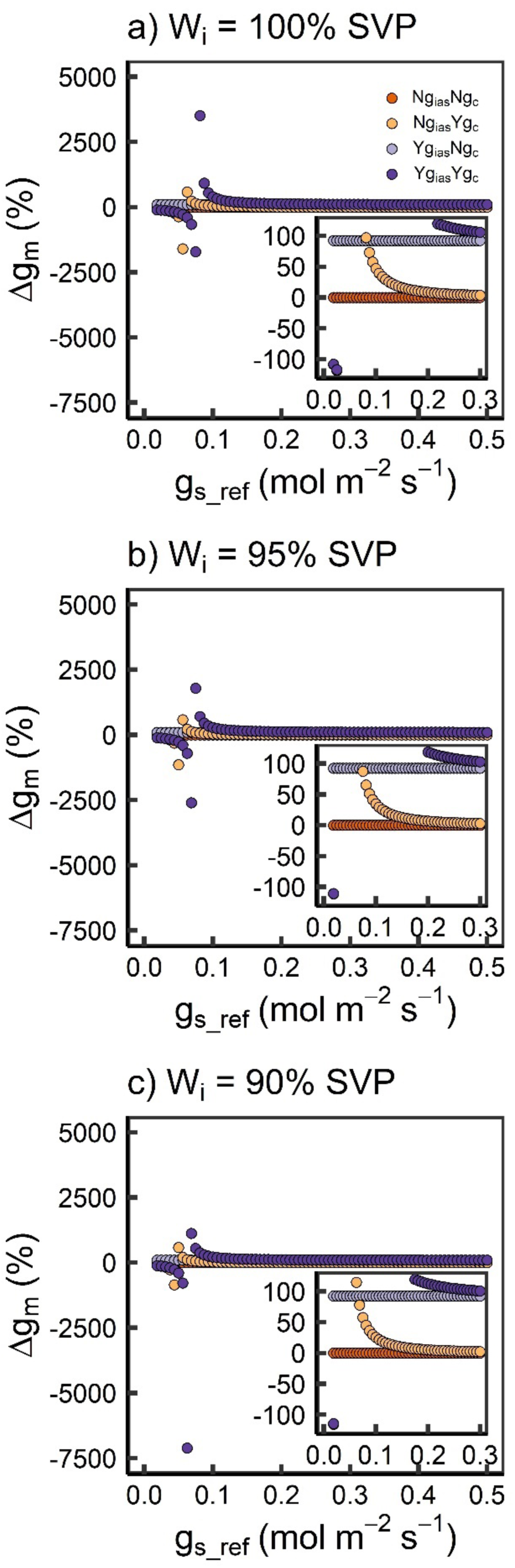
Modelled impacts of intercellular [H_2_O] (W_i_), cuticle conductance (g_c_), and intercellular airspace conductance (g_ias_) on mesophyll conductance (g_m_) expressed as a change in g_m_ (Δg_m_ (%)), across a range of reference g_s_ values (g_s_ref_). W_i_ is either a) 100% saturation vapor presure (e_s_), b) 95% e_s_, or c) 90% e_s_. Reference value for g_m_ is 0.48 mol m^-2^ s^-1^ at 1 atmosphere. For Δg_m_ < 10% in Ng_ias_Yg_c_, g_s_ref_ must exceed 0.187 mol m^-2^ s^-1^, 0.168 mol m^-2^ s^-1^, and 0.15 mol m^-2^ s^-1^ in a), b), and c), respectively. Ng_ias_: no CO_2_ gradient; Yg_ias_: CO_2_ gradient present, g_ias_ = 1000 mmol m^-2^ s^-1^; Ng_c_: g_c_ = 0 mmol m^-2^ s^-1^; Yg_c_: g_c_ = 10 mmol m^-2^ s^-1^.

So far, the above cases refer to conditions where W_i_ is at saturation vapor pressure. in cases where W_i_ is not at saturation vapor pressure inside the leaf (e.g. Cernusak et al., 2018), the consequences vary with the degree to which the assumption is violated. C_i_, g_sw_, and V_cmax_ are relatively unaffected when W_i_ is at 99% saturation (data not shown; note that xylem water potential is ∼ −200 kPa (Nonami & Boyer, 1987), which would have a W_i_ value of ∼99.85% saturation vapor pressure), however these parameters become both over- and under-estimated when W_i_ reaches 90% saturation depending on which assumptions are violated (Fig. 6). Thus, it appears that very small violations of this assumption will have minimal effects on gas exchange parameters. But caution must be exercised in cases where this assumption is likely to be violated, such as high vapor pressure deficit conditions, drought stress, and at high temperatures, as parameters will be overestimated. We note that there could be cases of multiple assumption violations leading to ‘correct’ parameters for the wrong reasons (i.e. Figs. 6h,i), although even in these cases other parameters are still different.

Based on our modeling, we predict that under conditions where g_sw_ is low (well-watered, low light, high [CO_2_], low vapor pressure deficit, high leaf water potential), cuticular water loss will be sufficient to cause calculations to overestimate C_i_ (Fig. 2). Under conditions where W_i_ is less than expected (drought, high vapor pressure deficit, high temperature), calculated C_i_ values will be lower than the actual C_i_ (inside the mesophyll cells in Fig. 2b). As g_ias_ becomes increasingly finite, the C_i_ estimates will change in meaning from C_ias_ to C_i,es_, barring violations in the other assumptions. Lastly, under conditions where g_sw_ is high, g_cw_ is minimal, g_ias_ is very high, and the assumptions of Moss and Rawlins (1963) hold, then calculated C_i_ and measured C_i_ should agree. However, in this last case, the agreement results from C_i_ meaning C_ias_, which means that the assumptions behind g_m_ measurements need to be adjusted accordingly.

Cuticular water loss could be a significant source of water flux across the leaf and has the potential to undermine the assumptions upon which gas exchange calculations are based. We urge caution when performing and interpreting measurements of g_m_ due to the potential impact of g_cw_ on g_m_ calculations. More research is needed to assess the magnitude of cuticular water loss across species and climates, however our data suggest that g_cw_ must exceed 1% of g_sw_ to have a substantial impact on photosynthetic gas exchange. Current information on the g_cw_ suggests that it may exceed that 1% threshold on average, depending on the measurement methodology. Combining the information from Riederer and Schreiber (2001) and Schuster et al. (2017), median g_c_ based on all methods was 1.84 mmol m^-2^ s^-1^ (IQR: 0.45 – 4.67 mmol m^-2^ s^-1^, mean: 2.73 mmol m^-2^ s^-1^), while median g_c_ based on permeance was 0.32 mmol m^-2^ s^-1^ (IQR: 0.11 – 0.90 mmol m^-2^ s^-1^, mean: 0.85 mmol m^-2^ s^-1^) and 3.31 mmol m^-2^ s^-1^ (IQR: 1.58 – 5.91 mmol m^-2^ s^-1^, mean: 3.82 mmol m^-2^ s^-1^) based on minimum conductance methods (Fig. 1a). Meanwhile, g_sw_ from the global datasets of Lin et al. (2015) and Smith and Dukes (2017) was 104 mmol m^-2^ s^-1^ (IQR: 52 – 226 mmol m^-2^ s^-1^, mean: 185 mmol m^-2^ s^-1^), suggesting that based on median values, cuticle conductance could range between and 3.2 % of g_sw_ (Fig. 1). Schuster et al. (2017) note that minimum conductance methods can be biased towards high values if stomata are not completely closed, which may explain the 10-fold difference in median permeance between the methods. Another possible explanation for this discrepancy could be related to stretching of the cuticle at high Ψ, which enhances g_cw_ (Boyer et al., 1997) and would be the case for many minimum conductance measurements but not for permeance methods, where isolated cuticle could shrink (Boyer, 2015). We also note the large discrepancy in sample sizes for estimates of g_sw_ (> 22,000 observations) and g_cw_ (404 observations and only 148 for permeance-based methods). The development of a rapid method for assessing g_cw_ would help in circumventing broken assumptions when g_cw_/g_sw_ is high.

The source of cuticular water loss is (un)clear: some evidence suggests that the bulk of cuticular water loss occurs across the guard cell cuticles rather than the epidermal surface (Šantrůček et al., 2004). Such heterogeneity in cuticular conductance across a leaf would need to be accounted for to obtain accurate C_i_ estimates, especially if epidermal and guard cell cuticular water loss show differential responses to leaf turgor. This would only matter if the change in permeability with turgor of the guard cells was greater than the change in permeability with turgor in other cells on the leaf surface. However, much of the work measuring cuticle conductance focuses on either isolated cuticle from astomatous leaf surface or a gravimetric determination of cuticle conductance, with assumptions on stomatal opening and closure.

However, we note that part of the apparent effect of cuticle conductance in some studies may be due to CO_2_ gradients (finite g_ias_) within leaves. In fact, both processes can result in similar effects at low g_sw_ (Parkhurst, 1994; Fig. 6a), and each process could explain the evidence supporting the other process making partitioning difficult. Therefore, partitioning the impacts of g_cw_ and g_ias_ on C_i_ estimates should be a research priority.

## Recommendations

We recommend the following:

1. Whenever possible, measure water potential (ideally Ψ_m,apo_) to estimate W_i_ inside the leaf.
2. If measurements are not possible, choose suitable values of Ψ from the literature and calculate W_i_.
3. Be clear as to the definition of C_i_: is it C_i,es_, C_ias_ or some other value? This will ensure that gas exchange parameters can be properly compared without confounding different aspects of leaf physiology.

In terms of understanding g_m_, it is apparent that splitting g_m_ into its component parts (e.g. g_ias_, g_liq_) is necessary to understand how internal conductances respond to the environment. Given the likelihood of the variable location of C_i_ during most g_m_ measurements, many of the g_m_ measurements may not be directly comparable as they would be comparing different resistance pathways. Given that recent data assessing W_i_ inside leaves focused on xerophytic leaves (Cernusak et al., 2018), more data are needed to understand how W_i_ varies across environmental conditions in more mesophytic species, and especially angiosperms.

As a community, we have made significant advances within the Moss & Rawlins (1963) paradigm. Technological advances are now making it possible and crucial to move beyond the Moss & Rawlins paradigm to further our understanding of photosynthesis and gaseous diffusion in leaves by addressing each of the assumptions (Fig. 8).

**Figure 8.**
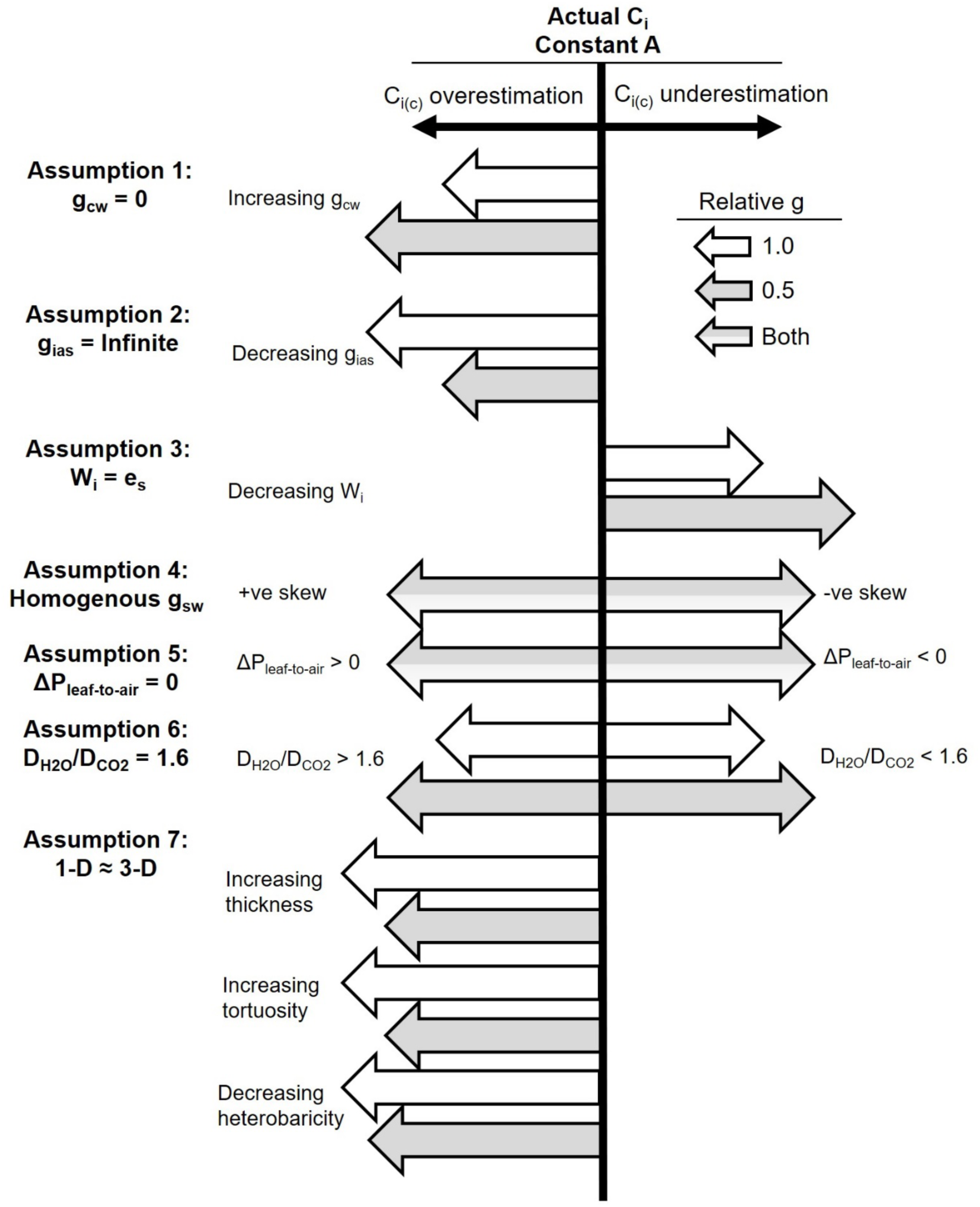
Relative effects of departures from each assumption on calculated C_i_ (C_ic_) relative to actual C_i_ under constant A and constant measured conductance (g). **Assumption 1**: increasing g_cw_ causes C_ic_ to increase, leading to overestimation of C_i_, with a stronger effect at lower g. **Assumption 2**: decreasing g_ias_ causes C_ic_ to increase, leading to overestimation of C_i_, with a stronger effect at higher g. **Assumption 3**: decreasing W_i_ from 100% SVP causes C_ic_ to decrease, leading to underestimation of C_i_, with a stronger effect at lower g. Decreasing leaf water potential (Ψ_m,apo_) causes W_i_ to decrease, causing C_ic_ to decrease and underestimate C_i_. **Assumption 4**: positively skewed stomatal apertures means that stomata are more closed than expected based on g, such that C_ic_ overestimates C_i_. Likewise, negatively skewed stomatal apertures means that stomata are more open than expected based on g, such that C_ic_ underestimates C_i_. This results from influences on g_ias_ – smaller than expected stomatal apertures means that the effective pathlength for diffusion is longer, decreasing g_ias_ relative to an “expected value”, while larger than expected stomatal apertures means that the effective pathlength for diffusion is smaller, increasing g_ias_ relative to an “expected value”. In the second case and under Assumption 2, the difference in g_ias_ would be a difference between a smaller and larger infinite value. **Assumption 5**: a pressurized leaf relative to air (ΔP_leaf-to-air_ > 0) means that calculated W_i_ is higher than the expected e_s_, causing C_ic_ to overestimate C_i_, and a negatively pressurized leaf (ΔP_leaf-to-air_ > 0) means that W_i_ is lower than the expected e_s_, causing C_ic_ to underestimate C_i_. Note that the pressure will also have implications for the diffusion dynamics, but we do not address them here. **Assumption 6**: when D_H2O_/D_CO2_ > 1.6, g_sc_ is overestimated, causing C_ic_ to overestimate C_i_, while D_H2O_/D_CO2_ < 1.6 causes underestimation of g_sc_, leading C_ic_ to underestimate C_i_. Note that these effects are stronger under smaller values of g. **Assumption 7**: when considering gas exchange in 3-D, tortuosity of the pathway and homo/heterobaricity of the leaf (which could feed into tortuosity), and leaf thickness impact g_ias_, with higher tortuosity, lower heterobaricity and thicker leaves, decreasing g_ias_, causing C_ic_ to overestimate Ci. **Note**: tortuosity, heterobaricity, and leaf thickness will influence g_ias_. Assumptions 3 and 5 feed into assumption 3, while assumptions 4 and 7 feed into assumption 2.

### Code and Data

All code is available as supplementary files (“Modeling.rmd”, “Reanalysis of gm data.rmd”, “Literature gc-gs-gias calculations.rmd”), as are input data (“popexample.csv”, “CO2response.csv”, “temp data.csv”, “gmlight.csv”, “cuticle conductance temp response.csv”, “cuticle_bins.csv”, “WP.csv”, “Diffusion bins Onoda data.csv”). Modeling code automatically generates .csv files for the modeling analysis.

## Acknowledgements

We would like to thank Dr. Patrick J. Hudson for comments on an early draft of the manuscript and Dr. John S. Boyer for comments on a late draft. We would like to thank Yusuke Onoda for providing data on leaf thicknesses from Onoda et al. (2011) and Jordi Martínez-Vilalta for providing water potential data from Martínez-Vilalta et al. (2014). This work was also supported by funding to DTH through the NSF EPSCoR Program under Award # IIA-1301346 and through NSF IOS 1658951 at the University of New Mexico. Any opinions, findings, and conclusions or recommendations expressed in this material are those of the authors and do not necessarily reflect the views of the National Science Foundation. J.T. is supported by Research Fellowships for Young Scientists from the Japan Society for the Promotion of Science (JSPS).

## Author Contributions

All authors contributed to the design of the study. JRS performed the modeling. JRS wrote the manuscript with input from all authors.

